# Comparative transcriptomics reveals an extracellular worm argonaute as an ancestral regulator of LTR retrotransposons

**DOI:** 10.1101/2025.09.22.674508

**Authors:** Isaac Martínez-Ugalde, Kyriaki Neophytou, Yenetzi Villagrana-Pacheco, Adriana Orrego Durañona, Lewis Stevens, Xiaochen Du, Rowan Bancroft, Jessica L Hall, Amy B Pedersen, Mark Blaxter, Amy H Buck, Cei Abreu-Goodger

**Author notes:** Correspondence should be addressed to Cei Abreu-Goodger.

## Abstract

Safeguarding the genome from non-self elements is essential for reproduction, development, and ageing. One of the major threats to genome integrity is Transposable Elements (TEs), which can be post-transcriptionally silenced through small RNAs (sRNAs) and argonaute proteins. Recent work suggests TE-derived sRNAs may also act as virulence factors in host–pathogen interactions. During infection, the intestinal parasite *Heligmosomoides bakeri* secretes a single argonaute protein (exWAGO) and a wide variety of TE-derived sRNAs. Although exWAGO is highly expressed, conserved, and secreted by parasitic nematodes, its function and sRNA guide preference remain unclear. Using comparative transcriptomics of the sRNAs bound to exWAGO within parasites of rodents, livestock and humans, and its orthologs in *C. elegans*, we found that exWAGO is capable of loading sRNAs produced from all classes of TEs in addition to some protein-coding and non-coding transcripts. However, our results suggest that the ancestral endogenous function of exWAGO was likely linked to LTR retrotransposon regulation. To understand how this relates to potential extracellular functions of exWAGO we also examined the sRNAs bound to exWAGO secreted by *H. bakeri* in both vesicular and non-vesicular forms. Extracellular exWAGO preferentially loads sRNA guides derived from non-autonomous and fragmented LTRs, suggesting the existence of adaptable reservoirs of regulatory sRNAs with potential roles in cross-species RNA communication. Together, our results show that exWAGO is part of an evolutionarily conserved pathway for LTR retrotransposon regulation, while preferentially utilising degenerated elements as sources of secreted sRNAs.

## Introduction

Although typically neutral (Arkhipova, 2018), transposable elements (TE) can have deleterious effects on their host genomes, such as gene disruption or ectopic recombination. Disregulated TE activity can also induce genome expansion, potentially leading to genomic instability. To offset this, host genomes have adapted different molecular mechanisms, including non-coding RNA pathways, to regulate TE activity (Van Lopik et al., 2023; Wolf et al., 2020; W. Zhou et al., 2020). The PIWI-interacting RNA (piRNA) pathway, for instance, is conserved across metazoans (Calcino et al., 2018) and plays a key role in TE silencing through sequence-specific targeting. In *Caenorhabditis elegans,* the PIWI argonaute (PRG-1) and its bound piRNAs trigger TE-derived silencing, inducing the amplification of antisense secondary small interfering RNAs (known as 22G siRNAs) by RNA-dependent RNA polymerases, which are loaded by worm-specific argonautes (WAGOs) (reviewed by Fischer 2024). These WAGO-22G complexes are ultimately responsible for TE silencing. Many clades of nematodes, however, have independently lost essential components of the PIWI pathway, including either the PIWI argonaute protein or the capacity to produce piRNAs (Beltran et al., 2019; Sarkies et al., 2015). The clade V of nematodes (divided into Strongylida and Rhabditina) have mostly maintained PIWI argonaute and the capacity to produce piRNAs. Strongylida parasites, however, show oversized genomes with large TE content, compared to the free-living Rhabditina (International Helminth Genomes Consortium, 2019). Despite the presence of the PIWI pathway in these nematodes, little is known about how TEs are regulated beyond *C. elegans*.

The gastrointestinal parasite of mice *Heligmosomoides bakeri,* secretes immunosuppressive extracellular vesicles (EVs) during infection, which carry TE-derived 22G siRNAs and the extracellular worm argonaute (exWAGO) (Buck et al., 2014; Chow et al., 2019). *H. bakeri*, is closely related to human infective nematodes (including the human hookworms *Necator americanus*, and *Ancylostoma* spp.), which collectively affect close to 25% of the human population (Stevens et al., 2023; World Health Organization, 2020). Given the limited anthelmintic options and the increase in resistance to them, novel therapeutic options are required. We recently showed that *H. bakeri* secretes exWAGO, both inside EVs and in a vesicle-free form, and that immunisation against exWAGO confers partial protection against *H. bakeri* infection (Neophytou et al. 2025). Both exWAGO forms associate with TE-derived small RNAs and can be internalized by host cells, suggesting a potential role in RNA-based cross-species communication (Neophytou et al., 2025). Furthermore, exWAGO is conserved among clade V nematodes and highly expressed relative to other argonaute proteins (Chow et al., 2019). Biochemical and proteomic analyses show its orthologs can be secreted by different species, including *C. elegans*, human-infective *Ancylostoma ceylanicum*, ruminant-infective *Teladorsagia circumcincta* and *Trichostrongylus colubriformis*, and rat-infective *Nippostrongylus brasiliensis* (Chow et al., 2019; Neophytou et al., 2025; Nikonorova et al., 2022; Rooney et al., 2022; Uzoechi et al., 2023). While the role of exWAGO in the extracellular environment is potentially linked to extracellular communication including parasite-host communication, its endogenous function and the features that drive its preference for loading specific sRNA guides remain uncharacterized.

Here, we use comparative genomics and transcriptomics of exWAGO-bound sRNAs in clade V nematodes to investigate how exWAGO guide preference evolved, and to probe how this underlies both genome defence and extracellular RNA export. Our findings suggest that the ancestral role of exWAGO was to regulate LTR retrotransposons. We also identify specific properties of secreted LTR-derived sRNAs in *H. bakeri,* highlighting how non-autonomous LTR fragments are a preferential source for potential cross-species RNA communication functions in parasitic nematodes.

## Results

### Improving transposable element annotation in Strongylida genomes

We previously observed that exWAGO prefers transposon-associated siRNAs (Chow et al., 2019; Neophytou et al., 2025). Yet to properly understand transposon regulation, we need high-quality annotations of genomic repeats. To get an evolutionary perspective on transposon regulation, we first improve the repeat annotations for the *H. bakeri*, *H. polygyrus*, *N. brasiliensis*, *T. circumcincta* and *A. ceylanicum* genomes, using *de novo* prediction followed by automatic and manual curation (see Methods: Transposable element annotation).

To assess if our curated repeat libraries improved the quality of genome annotations, we compared them against available repeat libraries from WormBase ParaSite and previous annotation projects (Chow et al., 2019; Howe et al., 2017). Given the lack of a previous repeat library, *H. polygyrus* was excluded from this comparison. We first compared the number and length of non-redundant consensus sequences of the most common TE categories: DNA transposons (DNA), long interspersed nuclear elements (LINEs), long terminal repeat retrotransposons (LTRs), short interspersed nuclear elements (SINEs) and Unknown repeats (Unknown). Our curated libraries in general contain a similar or increased number of consensus sequences for DNA, LINE and LTR elements, but a decrease in Unknown repeats, suggesting the latter have been incorporated into named classes (Fig. 1A). Our curated models also show a significant increase in length regardless of the category (one-tailed Wilcoxon rank-sum test, p < 0.01) (Fig. 1A). Finally, regardless of TE classification, curated models encode a larger number of relevant proteins (e.g. transposase, reverse transcriptase) (Supplementary Table S1). Our curated models are thus not only longer but will also enhance functional analyses of TEs in these organisms.

**Figure 1.**
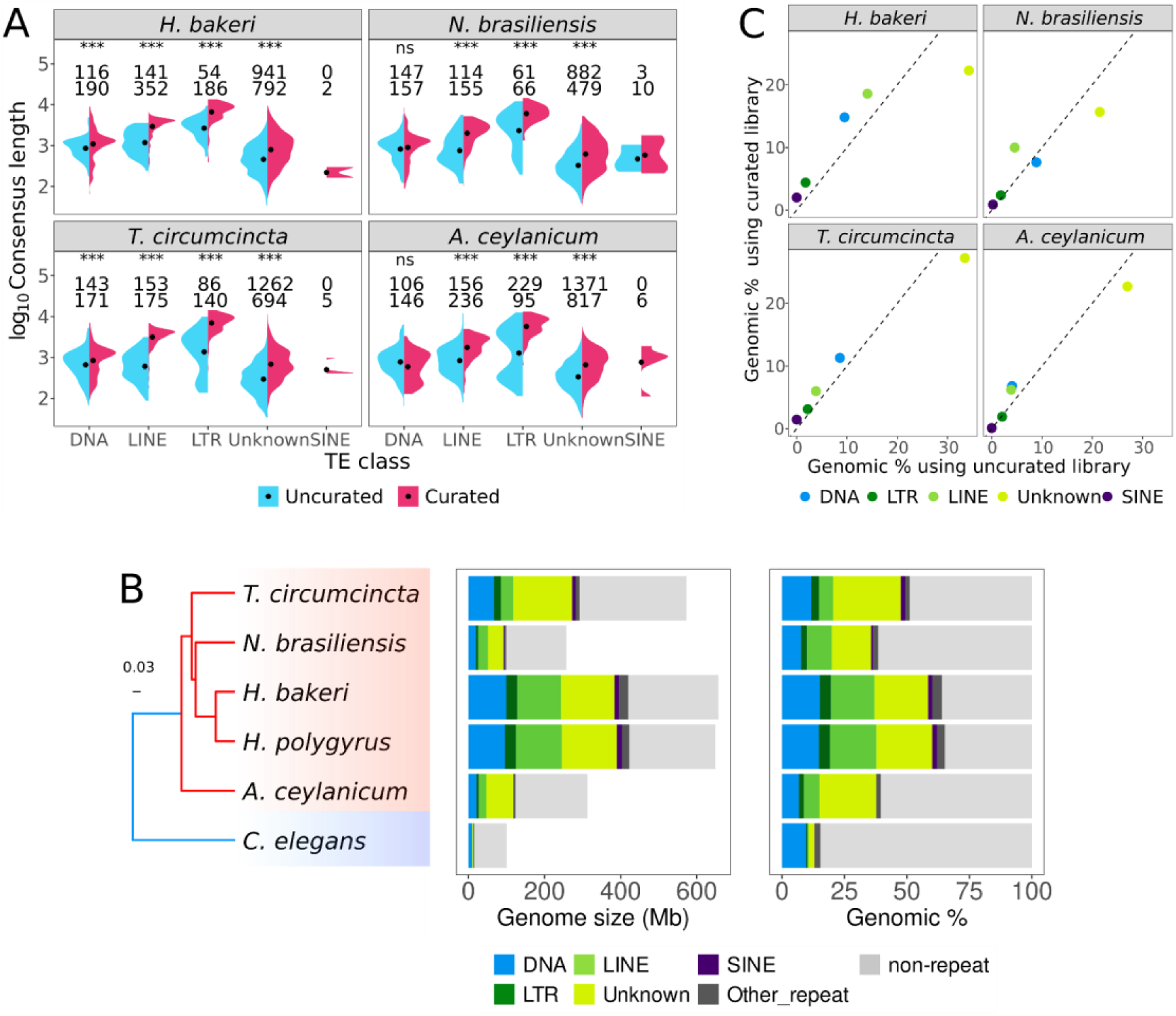
**Improved repeat annotations for Strongylida genomes**. Comparison of the number and length of uncurated and curated consensus models, by TE category (A). A one-tailed Wilcoxon Rank Sum test was used for comparisons (ns not-significant; * p-value < 0.05). The first/second row of numbers indicates the number of non-redundant uncurated/curated models. Phylogenetic relationship and repeat content in the analysed species based on a maximum likelihood phylogeny inferred with 443 BUSCO genes (parasitic species are highlighted in red and *C. elegans* in blue) (B). The repeat span is shown in megabases (Mb) and as a percentage of genome span (Genomic %). Comparison of the percentage of annotated bases per TE category between curated and uncurated libraries (C). The dotted line denotes x=y.

We further assessed the degree of similarity between uncurated and curated libraries using the benchmarking method provided by RepeatModeler2 (Flynn et al., 2020). Briefly, this benchmark uses a tested library to annotate a reference library, to then classify models based on their relative sequence similarity, coverage and divergence into perfect, good, present, and missed models. For *H. bakeri*, when comparing the uncurated against the curated library, we found 60 families with perfect agreement, 278 good, 509 present and 876 missing. This means that 876/1726 (51%) of the models in the curated library are missing in the uncurated library. We found similar proportions of missing models in the uncurated libraries for the rest of the species (mean= 39.8%, SD= 7.6%) (Supplementary Table S2). The number of missing families could reflect highly fragmented/divergent or genuinely absent families in the uncurated sets relative to the curated ones. Together, these results highlight important differences in the coverage and identity level between curated and uncurated libraries.

Our newly annotated genomes vary substantially in repeat content, consistent with their size differences, from 38% of the 257 Mb genome of *N. brasiliensis* to 65% of the 649 Mb genome of *H. bakeri* (Fig. 1B). Notably, the TE span in Strongylida is up to four times greater than the entire genome of *C. elegans* (100 Mb), underscoring substantial TE expansion in parasitic lineages.

Importantly, using our curated libraries, we substantially reduce the percentage of bases annotated as Unknown repeats in all the analysed species by 17-22% (Supplementary Table S3). For instance, in *H. bakeri,* Unknown repeats cover 34.2% of the genome (222 Mb) when using the uncurated library, while only 22.2% (144 Mb) when using the curated library (Fig. 1C). The reduction in bases annotated as Unknown is accompanied by an increase in all the other categories. For example, DNA transposons increase from 9.5% (61 Mb) to 14.7% (96 Mb), LINEs from 14% (91 Mb) to 18.5% (120 Mb) and LTRs from 1.7% (11 Mb) to 4.4% (28 Mb). Although we were able to decrease the span of Unknown repeats in the four species we compared (Supplementary Table S3), it is important to note that Unknown are still the most abundant category of repeats (Fig. 1C), highlighting the value of further curation and characterization efforts.

The improvement in the classification of repeats is particularly meaningful since we previously highlighted Unknown repeats as relevant contributors of exWAGO guides in *H. bakeri* (referred to as Novel repeats in Chow et al., 2019). By refining repeat annotations, we not only increase the accuracy of TE classification but also provide the foundation to improve our understanding of exWAGO guides.

### Improving the detection of exWAGO-bound sRNAs

To interrogate the sRNA guides bound to exWAGO in different Strongylida parasites, we used sRNA sequencing libraries generated following immunoprecipitation of exWAGO from adult worms. ExWAGO was pulled-down using rat polyclonal anti-exWAGO antibodies raised against recombinant *H. bakeri* exWAGO (for *H. bakeri*, *H. polygyrus*, *N. brasiliensis* and *A. ceylanicum*) or recombinant *T. circumcincta* exWAGO (for *T. circumcincta*) (Neophytou et al., 2025). Western blot analyses show that immunoprecipitations (IP) were successful, but pull-down efficiency varied for the different species (Neophytou et al., 2025). We generated sRNA libraries from the eluate of the IP and total worm lysates using RNA 5’ polyphosphatase treatment to allow capture of 5’ triphosphorylated sRNAs (Neophytou et al., 2025). To be able to compare against the binding preference of a free-living rhabditid species, we used published IP data for the exWAGO orthologs in *C. elegans*: SAGO-1, SAGO-2 and PPW-1 (Seroussi et al., 2023).

Sequenced reads from IP experiments represent a subset of the total sRNA population in adult worms (Adult total), showing similar length distribution and genomic origin (Fig. 2A/B). To determine which sRNAs are enriched in exWAGO, we quantified the sRNAs mapping to each annotated genomic region for exWAGO-IP and libraries from adult worm lysates. We then performed a differential expression analysis, comparing IP to Adult total (see Methods). Regions with significantly more sRNAs in the IP, which we call IP-enriched, should represent regions that produce siRNAs specifically bound by exWAGO, while regions with similar amounts of sRNAs in the IP and the input include unbound RNA fragments and those that are non-specifically bound. Initially, we used edgeR with default Trimmed Mean of M-values normalisation (TMM). With this approach, we found 612, 1,591, 978, 1,006, and 3,488 IP-enriched regions for *H. bakeri*, *H. polygyrus*, *N. brasiliensis*, *T. circumcincta*, and *A. ceylanicum*, respectively. For *C. elegans* we found 4,216, 13,363 and 1,413 IP-enriched regions for SAGO-1, SAGO-2 and PPW-1, respectively. However, the log_2_ Fold-Change (logFC) values show a bimodal distribution, the higher mode likely containing exWAGO-bound sRNA and the lower mode likely representing unbound sRNA (Fig. 2C and Fig. S1A). The default normalisation (TMM) is based on the hypothesis that most regions do not change expression, and in this case centres the higher mode (the likely exWAGO-bound sRNAs) on logFC=zero, since it contains more regions than the lower mode. This suggests that “most regions” are actually producing exWAGO-bound sRNAs, with the unbound regions representing a relative minority. Hence, the default normalisation method (TMM) cancels this biological effect, thereby hindering the detection of true argonaute-associated sRNAs.

**Figure 2.**
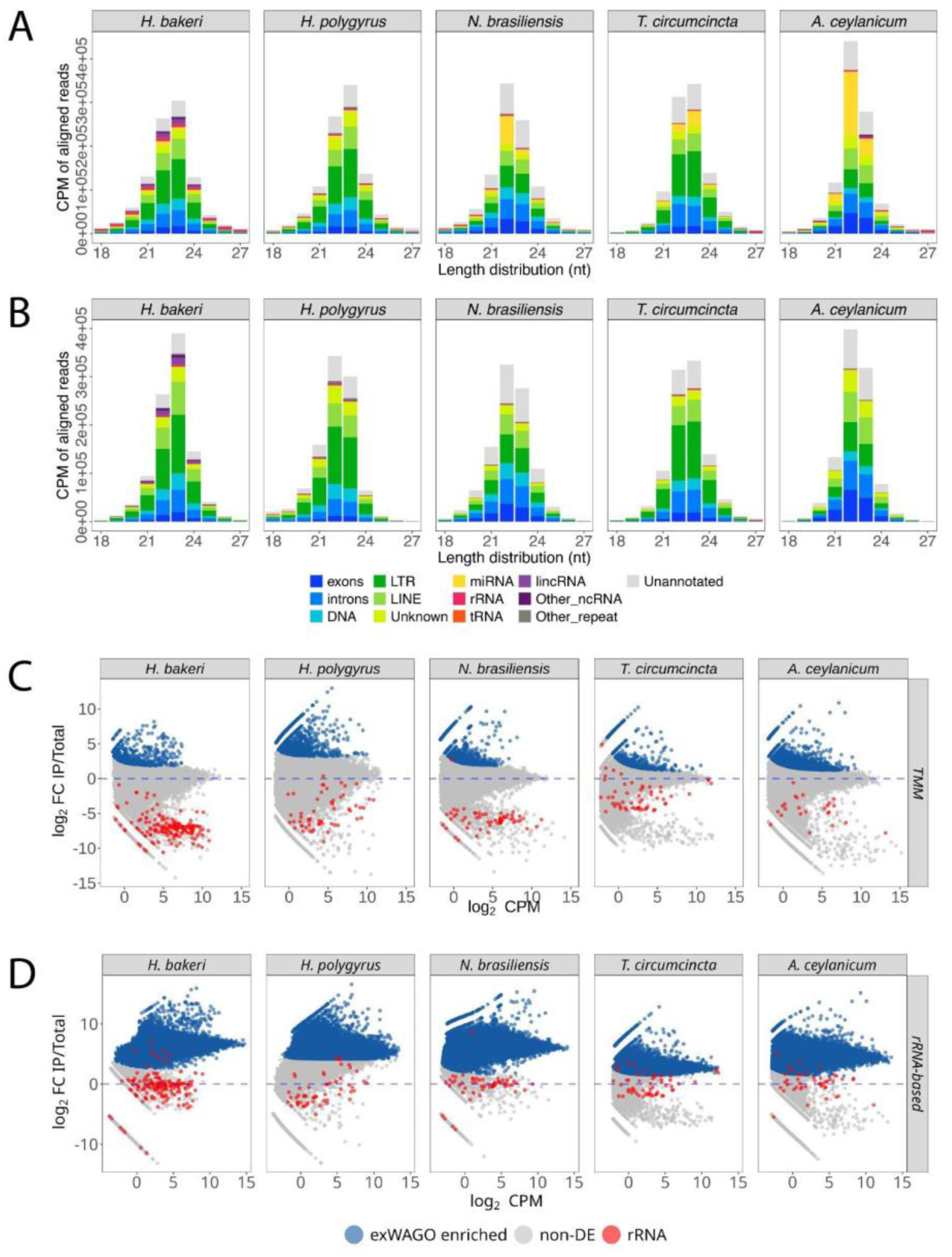
Analysis of exWAGO guides. Length distribution of aligned reads by annotation category for Adult total (A) or exWAGO IP (B) libraries in Strongylida parasites. The x-axis shows the length of aligned sRNAs (18-27 nt), while the y-axis shows average counts per million (CPM) across replicate libraries. MA plots (logCPM vs logFC) after differential expression analysis of exWAGO IP against total libraries using TMM (C) or housekeeping (D) normalisation. Each dot represents a non-overlapping region in each genome. Genomic regions identified as significantly higher in exWAGO IP vs Adult total (FDR < 0.01, see Methods) and interpreted as exWAGO-bound are shown in blue, while non-deferentially expressed regions (non-DE) interpreted as exWAGO-unbound are shown in grey. Regions annotated as rRNA are indicated in red. Dashed lines indicate logFC = zero in each contrast.

To address this problem, we turned to a “housekeeping” normalisation approach (Bullard et al., 2010). This relies on estimating normalisation factors using a set of housekeeping genes (those expected not to change in expression) so that the logFC values of this subset of genes are centred on zero. In our case, we had to choose sRNAs that are expected not to be bound by argonaute proteins. Thus, we used the sRNAs mapping to the sense-strand of rRNA genes to estimate normalisation factors (see Methods: Differential expression and enrichment analyses of exWAGO and adult total across clade V nematodes). We assume that these rRNA fragments are likely degradation products, and we expect most of them not to be specifically bound by argonaute proteins (Seroussi et al., 2023; Wynant et al., 2017; Zagoskin et al., 2022). To allow for the possibility that some rRNA fragments might be specifically bound and to reduce the effect of outliers, we applied the TMM principle to the rRNA counts to estimate more stable normalisation factors.

After applying the new normalisation, rRNA genes are consistently centred around logFC=0 (Fig. 2D), and we detect 148,185, 96,605, 140,238, 189,676 and 133,022 exWAGO-bound regions as being significantly enriched for *H. bakeri*, *H. polygyrus*, *N. brasiliensis*, *T. circumcincta,* and *A. ceylanicum*, respectively. These results suggest that in Strongylida, exWAGO binds sRNAs from a large number of genomic regions. Importantly, the regions that we determine not to be bound by exWAGO show a shift towards miRNA and other ncRNA after housekeeping normalisation (Fig S2). Nonetheless, it is important to notice that even after using rRNA to estimate normalisation factors, some of the rRNA genes do fall within the exWAGO-bound regions, which could represent real rRNA-derived guides or reflect technical issues due to higher background and/or differing IP efficiency among samples.

Interestingly, when applying the same normalisation to the *C. elegans* orthologs of exWAGO, we do not see as substantial a change in enriched regions for SAGO-1 (4,216 to 3,943) or SAGO-2 (13,363 to 14,482), but for PPW-1 the number of enriched regions increases from 1,413 to 42,236, similar to the increase for exWAGO. We further analyzed other *C. elegans* argonaute (AGO) IP data (Seroussi et al., 2023), and we observed a large increase in the number of enriched regions for a few WAGOs, including WAGO-1 and PPW-2 (58,357 and 38,286, respectively), but not for other AGOs such as PRG-1 or ERGO-1 (Fig. S3 and S4). These results align with the expected behaviour of TMM, which centers the expression values where the majority of regions are distributed (Bullard et al., 2010). Therefore, in AGOs that mostly bind to specific types of sRNA, like ALG-1 or PRG-1, which bind miRNAs or piRNAs, respectively, most of the regions will be unbound. In contrast, AGOs which bind sRNAs from a broader number of genomic regions, as often occurs for siRNAs, we expect an increase in enriched regions when applying rRNA-based normalisation.

Finally, to check that our new analyses were not biased towards an increase in regions while sacrificing the quality of these, we quantified the sRNA abundance of other ncRNA species. We found that most regions annotated as ncRNA family species, such as tRNA, miRNA, snRNA, or snoRNA, remain unbound by exWAGO (Supplementary Table S4). On average, 69.8% of tRNAs remain unbound (SD = 24.4%), 86.1% of miRNAs (SD= 12.2%), 88.1% of snRNA (SD = 2.6%), and 93.5% of snoRNAs (SD = 7.9%). These data indicate that our rRNA-based normalisation results maintain biologically meaningful patterns, since WAGOs in general are not expected to bind these ncRNAs (Seroussi et al., 2023).

Taken together, our results indicate that exWAGO can bind sRNAs from a wide variety of genomic regions. This is consistent with what is observed for other WAGO proteins in *C. elegans* (Seroussi et al., 2023).

### ExWAGO as a global regulator of transposable elements

Since exWAGO-bound sRNAs map to a large number of regions in each genome, we next asked if exWAGO shows specificity for certain types of annotation. For both *Heligmosomoides* species and *T. circumcincta*, 62-71% of the exWAGO-bound sRNA counts are derived from repetitive elements, while for *A. ceylanicum* and *N. brasiliensis,* this value is just under 48% (Fig. 3A). In *C. elegans,* this number is further reduced, where out of the three orthologs, only PPW-1 shows more than 14% IP-enriched sRNAs coming from repeats. The opposite trend is observed for protein-coding regions, where the *C. elegans* orthologs show 38-68% of sRNA counts, while in *Strongylida* only 3-12% of their exWAGO-bound sRNAs are derived from protein-coding regions. We further categorised sRNAs based on whether they mapped to the sense or antisense strand of each annotation. Antisense strand-derived sRNAs that map to exons, DNA transposons, LTR retrotransposons, LINEs and lincRNAs, are consistently more abundant than those derived from the sense strand (Fig. S5). This is consistent with WAGO guides being produced by RNA-dependent RNA polymerases RdRps using a primary RNA as a template (Pak & Fire, 2007; Sarkies et al., 2015).

**Figure 3.**
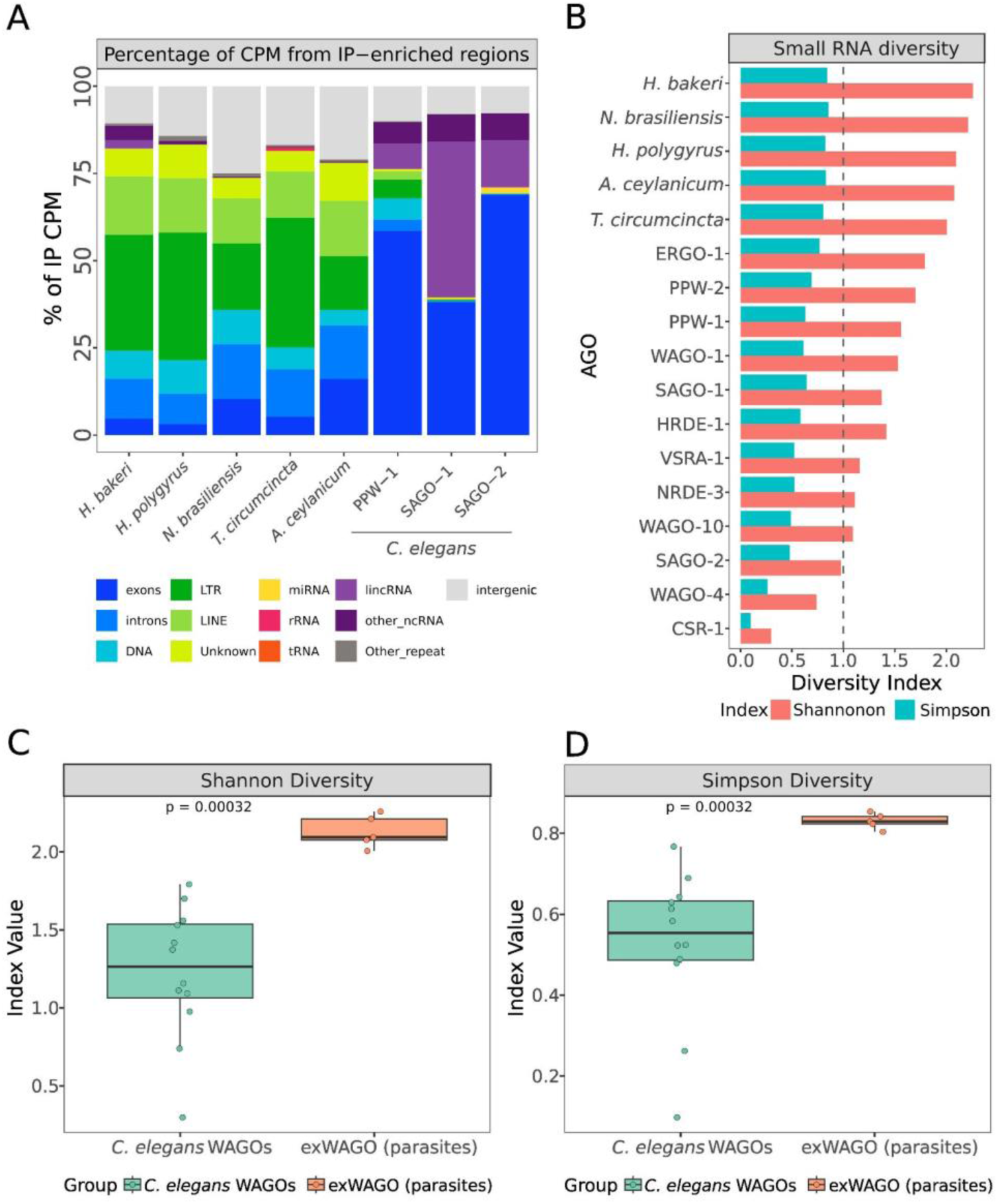
**Small RNA diversity in *C. elegans* WAGOs and exWAGO in Strongylida parasites**. A comparison of the percentage of Counts Per Million (CPM) of enriched regions per genomic category after differential expression analysis is shown in (A). Small RNA diversity represented by Shanon and Simpson indices in *C. elegans* WAGOs and exWAGO in Strongylida parasites (B). The grey dotted line indicates the maximum value (1) of the Simpson index. Box plots comparing the distribution of Shannon and Simpson indices between *C. elegans* WAGOs and exWAGO in Strongylida parasites (C-D). Panels B–D are based on the average of 100 rarefied random samplings using the number of enriched regions in WAGO-10 (2,029 regions) as the standardized sampling size.

Notably, in Strongylida parasites, 98-100% of the TE families with detectable sRNAs in the Adult libraries have at least one copy enriched in exWAGO IP. This includes DNA transposons, LTRs, LINEs, SINEs and Unknown repeats, suggesting that exWAGO could regulate most of the expressed TEs in these parasites. Similarly, the *C. elegans* ortholog PPW-1 binds sRNAs from 99% of the TE families with detectable sRNA production. Even though SAGO-1 and SAGO-2 have a shared evolutionary history with PPW-1, they only bind sRNAs for a reduced set of TE families with detectable sRNAs (15.7% and 16.2%, respectively), suggesting subfunctionalization of these WAGOs, possibly due to differences in tissue localization. Notably, whilst SAGO-1 and SAGO-2 are mainly expressed at the apical membrane of the intestine, PPW-1 shows enrichment for both germline and the apical membrane of the intestine (Seroussi et al., 2023).

To gain more insight into the diversity of exWAGO guides, we expanded our analysis to other WAGOs in *C. elegans* (Seroussi et al., 2023), where we calculated Shannon entropy and Simpson’s diversity indices using a comparable set of 43 genomic features, including exons, introns, DNA transposons, LTRs, LINEs, and different ncRNA families (Supplementary Table S5). We normalized counts using average counts per million (CPM) to account for differences in sequencing depth. In this context, Shannon entropy reflects the evenness in sRNA guide production across different annotation categories, while Simpson’s diversity highlights the dominance of specific categories. Highly specialized AGOs such as ALG-1 or PRG-1 show low Shannon entropy (0.004 and 0.3, respectively) and low Simpson diversity (0.001 and 0.11) (see Fig. S6), highlighting low diversity and dominance of specific categories. This is expected since ALG-1 has a high selectivity for miRNAs, and PRG-1 for piRNAs. However, Shannon entropy in *C. elegans* WAGOs ranges from 0.3 to 2.02 (mean=1.3, SD=0.55), whilst Simpson’s diversity ranges from 0.09 to 0.79 (mean=0.1, SD=0.79).

Notably, exWAGO in Strongylida exhibit a significantly higher diversity in comparison to *C. elegans*’ AGOs (Wilcoxon one-tailed rank-sum test, p-value < 0.001) in both Shannon (mean=2.22, SD=0.11) and Simpson indices (mean=0.84, SD=0.02). This comparison, however, could be biased by the number of regions producing enriched sRNA in these AGO proteins, which is directly related to genome size. Therefore, to account for these differences, we randomly sub-sample the data for each genome to match the smallest number of enriched regions (WAGO-10, 2,029 genomic regions), but only considering WAGOs, due to the high selectivity of other AGOs like ALGs or PRG-1. For this, we randomly sampled 100 times each set of enriched regions (Fig. 3B). Even using this reduced set, average Shannon and Simpson indices confirm that exWAGO in Strongylida is significantly more diverse than *C. elegans* WAGOs (Wilcoxon one-tailed rank-sum test, p-value < 0.001) (Fig. 3C and D).

Together, these results suggest that the ancestral exWAGO protein was a versatile argonaute, capable of loading sRNAs derived from a wide variety of sources, including most transposon families present in the ancestral genome as well as a subset of protein-coding and non-coding transcripts. This hypothesis is supported by the conserved capacity of PPW-1 to load TE-derived sRNAs, despite the comparatively reduced diversity (likely due to specialisation) of its paralogs SAGO-1 and SAGO-2. The strong relationship between exWAGO and TE-derived sRNAs, coupled with its diversity in Strongylida, raises the possibility that this AGO protein has been critical for regulating genome stability in species with high transposable element (TE) activity, ultimately shaping genome architecture over a broad evolutionary timescale.

### ExWAGO as an ancestral regulator of LTR retrotransposons

Although exWAGO binds to a highly diverse set of sRNAs in all parasites, this does not rule out a preference for certain sRNA classes over others. To account for the effect of differences in genome size and annotation coverage, we calculated the density of sRNA for each annotation category (see Methods). Although LTRs are not the most abundant repeats (Fig. 4A), antisense LTR-derived sRNAs show the highest density across Strongylida with 6.2-12.6-fold enrichment relative to their genomic span (Fig. 4B). Antisense exon-derived sRNAs are the second highest category, with an average 3.2-fold enrichment in parasites. In contrast, PPW-1 in *C. elegans* shows a preference for antisense lincRNA-derived sRNAs (7.3-fold enrichment), followed by antisense LTR-derived sRNAs (4.6-fold enrichment), with antisense exon-derived sRNAs in third place (3.1-fold enrichment). Interestingly, when antisense pseudogene-derived sRNAs are taken into account (a category we only have for *C. elegans*), they show a similar enrichment as antisense LTR-derived sRNAs (4.8-fold enrichment) (Fig S7). Consistent with previous reports (Seroussi et al., 2023), SAGO-1 and SAGO-2 show a strong preference for antisense lincRNA-derived sRNAs, with a 40.3-and 12.7-fold enrichment, respectively, suggesting the function of these orthologs has diverged from that of PPW-1.

**Figure 4.**
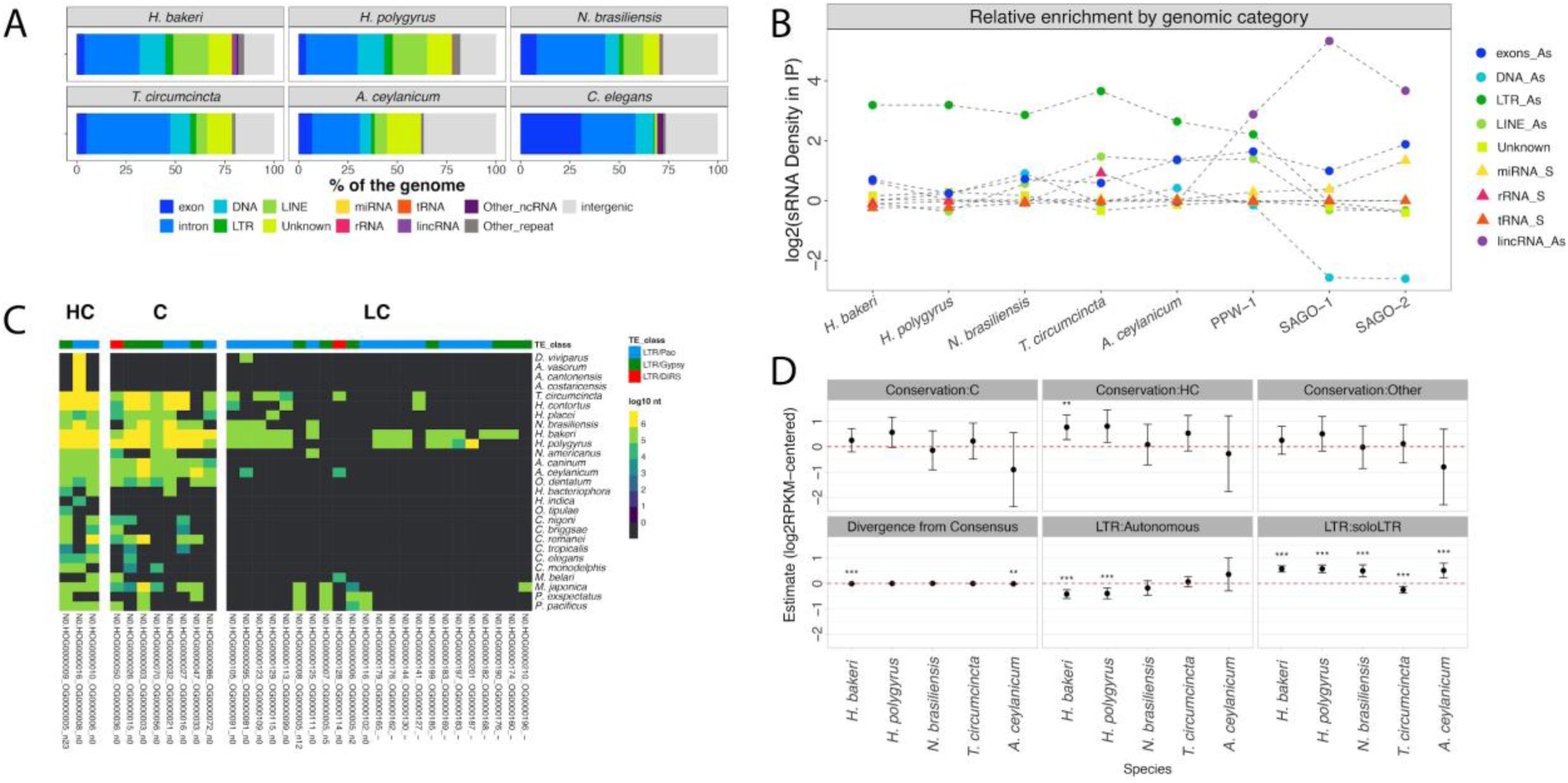
**LTR enrichment in exWAGO guide production**. Comparison of genome annotations in Strongylida parasites and *C. elegans* (A). Comparison of sRNA density of genomic regions enriched in exWAGO guide production per genomic category (B). The X-axis represents each species, while the Y-axis shows density values estimated as the log_2_ of the ratio of the percentage of CPM of IP-enriched regions and the percentage of genomic bases annotated for each of the categories. Phylogenetic orthology inference analysis of LTRs based on the reverse transcriptase (RT) protein (C). The top colour bar indicates LTR classification at the superfamily level. The heatmap shows the genomic abundance of the LTR in log_10_ of bases per HOG. Estimates for the fixed effects from the generalized linear mixed model (D). The red dashed line indicates 0 or a neutral effect (* p-value < 0.05, ** p-value < 0.01 and p-value < 0.001).

Certain genomic categories, especially lincRNAs and pseudogenes, are not easy to annotate *de novo*, leading them to be underrepresented or absent in genomes other than *C. elegans.* Nevertheless, our results suggest that the common ancestral function of exWAGO is related to LTR retrotransposon regulation, while some protein-coding genes also appear to be regulated in all species. This finding is consistent with previous observations in *H. bakeri*, indicating that the most abundant population of antisense siRNAs at the adult stage are retrotransposon-derived sRNAs and protein-coding-derived sRNAs (Chow et al., 2019). The consistent enrichment of LTR-derived sRNA guides across all species raises the question of whether highly conserved LTRs families, or more recently expanded (lineage-specific families), are the preferred source of exWAGO guides.

### Not conservation but structure drives LTR-derived exWAGO guide production

While some of the families involved in exWAGO guide production may be conserved across species, their capacity for transposition and degree of sequence divergence may also influence the levels of produced sRNAs. To assess the influence of evolutionary conservation of TE families on sRNA levels, we first annotated repetitive elements in 21 additional Strongylida and Rhabditina species encoding exWAGO using automatically curated libraries (Fig. S8). As observed in *H. bakeri*, *H. polygyrus*, *N. brasiliensis*, *T. circumcincta*, *A. ceylanicum,* and *C. elegans*, LTR retrotransposons are one of the repeat categories with the smallest genomic span across clade V nematodes. We then annotated LTR-encoded proteins and inferred orthology relationships across all identified families using TEsorter (along with HMM models from Rexdb) and OrthoFinder. Using predicted reverse transcriptase (RT) sequences, we defined phylogenetic hierarchical orthogroups (HOGs) and classified LTR families into four categories: highly conserved (present in ≥15 species), conserved (5–14 species), lowly conserved (<5 species), and undetermined (lacking a predicted RT) (Fig. 4C). To quantify sequence divergence, we calculated the relative distance between each LTR copy and its consensus using RepeatMasker. To assess structural completeness, we classified LTRs into three categories based on coding capacity: autonomous (encoding all transposition-related proteins), non-autonomous (missing some domains), and soloLTRs (lacking internal coding regions). Finally, to account for differences in element size due to coding capacity or divergence, we normalized sRNA abundance by copy length and library size using reads per kilobase per million (RPKM).

Having classified LTR retrotransposons by conservation, structural variation and estimated their divergence, we fitted a generalized linear mixed model. We used conservation (based on HOGs) and structural completeness (based on the presence of transposition-related proteins) as categorical fixed effects, while divergence as a continuous fixed effect. To account for the variation that different families can have due to differences in activity, we used family as a random effect. Although we initially fitted strand as a random effect, diagnostic plots show that this variable introduces systematic bias into the model, suggesting a confounding effect. Since there is a strong bias towards antisense sRNAs among exWAGO guides (Fig. S5), we used the log_2_ mean-centred RPKM of antisense LTRs as the response variable.

After assessing model fit across species—finding no major deviations from normality (Fig. S9)— we observed that, on average, the strongest and most consistent positive effect on sRNA production was associated with soloLTRs (relative to non-autonomous; mean β = 0.42 and SE = 0.09) (Fig. 4D) (Supplementary Table S6). In contrast, autonomous LTR retrotransposons only show a positive but non-significant effect in *A. ceylanicum* (β= 0.35, SE= 0.33, p-value = 0.27). Notably, the preference for soloLTR-derived sRNAs remains consistent even when compared separately to either the internal region or the LTR region of full-length LTR retrotransposons (Fig. S10). Meanwhile, high conservation shows a positive effect on sRNA production in some species (compared to lowly-conserved; β = 0.39, SE = 0.42, on average), however, there is greater uncertainty around this estimate, and in general, the effect is not significant. Finally, the effect of divergence seems to be modest (β = −0.003, SE = 0.002, on average) and negatively related to sRNA production, suggesting that less divergent copies (more recent insertions) produce slightly more sRNAs.

Together, our results suggest that the ancestral function of exWAGO in clade V nematodes is related to LTR retrotransposon regulation. Furthermore, despite being non-autonomous, soloLTRs are relevant for exWAGO guide production, suggesting that retaining these genomic remnants may represent an adaptive advantage either for enhancing TE regulation in *trans* to silence active elements of the same family or potential alternative functions of exWAGO. Notably, our results also suggest that similar to the exWAGO protein, the sources of sRNAs are also ancestrally conserved. These results indicate that exWAGO has been adapted to regulate a set of commonly active LTRs across clade V nematodes.

### Structural differences among LTRs associated with sRNA secretion pathways

We recently showed that exWAGO can be secreted by *H. bakeri* in two forms (inside and outside of EVs) and that exWAGO is internalized by host cells *in vivo* (Neophytou et al., 2025). Given that EVs containing exWAGO can induce host immunosuppression (Buck et al., 2014), and non-vesicular exWAGO can also enter host cells (Neophytou et al., 2025), we sought to identify the structural features of LTRs that influence the secretion of exWAGO-bound sRNAs. To tackle this, we used vesicular and non-vesicular exWAGO IP libraries from secreted materials.

We performed differential expression analyses to compare the vesicular or non-vesicular exWAGO libraries against the adult IP libraries (see Methods), including the structural-based classification of LTR retrotransposons. Using this approach, we identified a set of 17,950 and 22,548 genomic regions enriched in vesicular and non-vesicular exWAGO, respectively (Fig. 5A and 5B).

**Figure 5.**
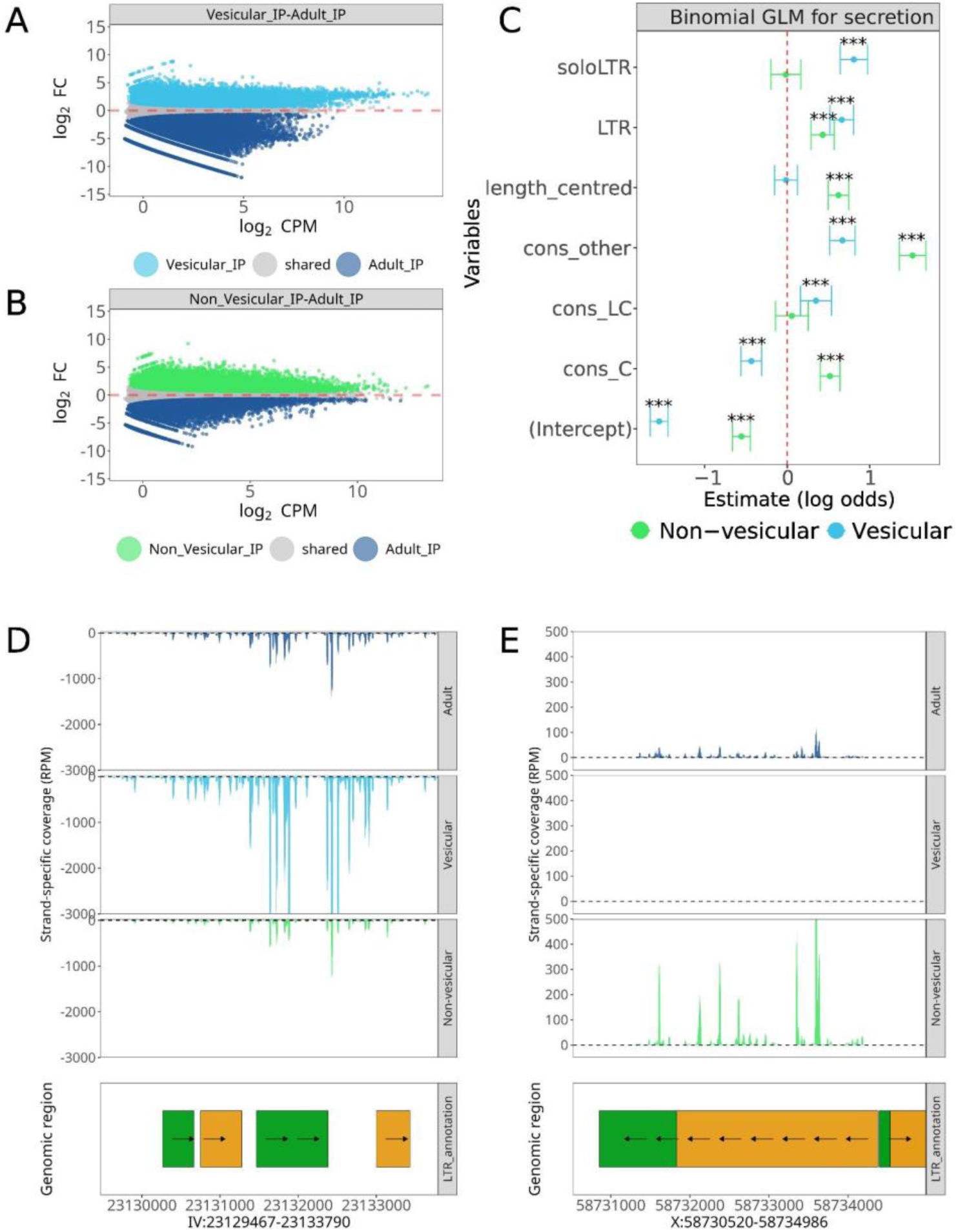
**Vesicular and non-vesicular exWAGO guides derive from different types of non-autonomous LTR retrotransposons**. MA plots (logCPM vs logFC) after differential expression analysis (FDR < 0.01) of vesicular exWAGO IP against exWAGO IP in adult worms (A), and non-vesicular exWAGO IP against Adult IP (B). The red dotted line indicates log_2_FC= zero. Estimated coefficients for binomial generalized linear models testing exWAGO LTR-derived small RNA secretion in vesicular and non-vesicular forms (C). Examples of vesicular (D) and non-vesicular (E) enriched regions showing sRNA production in Adult exWAGO IP (dark blue), vesicular exWAGO IP (light blue) and non-vesicular exWAGO IP (green). Genomic region tracks show LTR annotation (green for the LTR region, yellow for the internal coding region).

To determine if LTR retrotransposons involved in secreted exWAGO guide production differ from those that produce sRNA guides that remain inside the worm, we fitted binomial generalised linear models using the preference for secreted or retained exWAGO guide production as the response variable in the vesicular or non-vesicular contrast against adult, encoding regions enriched in secreted exWAGO as 1 and the remaining regions as 0. As explanatory variables, we used the previously defined degree of family conservation and structural classification. Moreover, we assessed length (centred on the average log_10_ length) to account for differences between different types of LTR retrotransposon structures. There were no significant deviations of model assumptions in simulated residuals for both models (Fig. S10 A and B).

The results of our model, for the vesicular exWAGO form, indicate that the odds of sRNA secretion increase on average 2.24 times when sRNAs are produced from soloLTR (β=0.80, SE=0.08) and 1.92 from the LTR region (β=0.66, SE= 0.07), relative to the internal coding region of an LTR, while holding the rest of the variables constant at their base levels (p < 0.001) (Fig. 5C and Supplementary Table S7). Moreover, we found that the odds of sRNA secretion from lowly conserved families (β=0.34, SE=0.09) or undetermined conservation (β=0.66, SE=0.07) due to the lack of detectable RT, significantly increase on average 1.42 and 1.95 times, respectively, compared to highly conserved families (p < 0.01). Notably, the mean-centred log_10_ length has a non-significant and negative effect (β=-0.01, SE=0.07, p-value < 0.84), supporting the preference for localised short regions of LTR retrotransposons, such as the LTR region or soloLTRs, as sources for vesicular-exWAGO guide production.

Our results for the non-vesicular form show that the odds of secretion increase 1.54 times when sRNAs are produced from LTR regions (β = 0.43, SE = 0.07, p < 0.001). Nevertheless, this preference is not from soloLTRs as with vesicles, but from longer albeit incomplete copies (Fig. 5C and Supplementary Table S8). Regarding conservation, families annotated as conserved (β = 0.52, SE = 0.06) and undetermined (β = 1.52, SE = 0.08) are 1.68 and 4.56 times more likely (p < 0.001 in both cases), compared to highly conserved families, whereas lowly conserved families show no significant association. Unexpectedly, we also found that length is positively associated with non-vesicular secretion (β = 0.62, SE = 0.06), corresponding to a 1.86 increase in odds for every 10-fold increase in length. The overall results indicate that while the vesicular exWAGO prefers sRNAs from soloLTRs, the non-vesicular form associates sRNAs from longer non-autonomous elements. Consistently, both vesicular and non-vesicular exWAGO are associated with LTR families with undetermined conservation (conservation Other), due to the lack of RT (Fig. 5C). Therefore, the lack of autonomy for retrotransposition increases the odds for serving as a source for secreted exWAGO guides.

To exemplify regions which produce significantly more secreted exWAGO guides, we highlight a soloLTR on chromosome IV with higher sRNA production (on average 14.44 log_2_ CPM) for the vesicular form (Fig. 5D), and a fragmented LTR retrotransposon on chromosome X (on average 11.8 log_2_ CPM) for the non-vesicular form (Fig. 5E). Both regions show clear differences in LTR structures, which are consistent with the overall preferences we found using binomial generalized linear models, with the vesicular form showing a preference for soloLTR-derived sRNAs, and the non-vesicular form for longer but non-autonomous elements.

Collectively, our results delineate two selective routes for the source of secreted exWAGO guides: vesicles enriched in lowly conserved, and degenerate soloLTRs, while the non-vesicular form prefers longer elements with intermediate conservation. Notably, in both secreted exWAGO forms, LTR-derived sRNAs are produced from non-autonomous elements, suggesting that once LTR retrotransposons lose their autonomous transposition capacity, they can be co-opted to act as the source of secreted sRNAs.

## Discussion

To mitigate the potential deleterious effects of TE activity, organisms have adapted molecular components, including argonaute proteins and their sRNA guides to recognize and silence TEs. Most of what we know about TE regulation in nematodes stems from their detailed annotation and functional characterization in *C. elegans*; however, much less is known about these elements and their regulation in parasitic nematodes.

Here, using manually curated TE annotations in multiple Strongylida parasites, we characterize the sRNA preferences of the extracellular worm argonaute exWAGO in an evolutionary context. Our improved TE annotations, combined with refined differential expression analysis, revealed exWAGO as a highly versatile AGO, but with an ancestral function linked to LTR retrotransposon regulation. Although exWAGO guides in adult Strongylida parasites preferentially originate from non-autonomous soloLTRs, the conservation of their corresponding TE family does not appear to influence the levels of sRNAs produced.

Interestingly, exWAGO preference for soloLTR-derived sRNAs extends to extracellular vesicles (EVs) in *H. bakeri*. However, secreted non-vesicular exWAGO shows a particular preference for sRNAs derived from longer and less degenerate elements. The structural differences between LTRs related to the two secreted exWAGO forms shape the composition of sRNAs delivered to the parasite’s host. The recurrent lack of detectable reverse transcriptase –and, therefore, autonomy for retrotransposition–of these elements indicates a consistent pattern of prior decay followed by co-option as a source of secreted sRNAs.

### Addressing compositional bias in Argonaute IPs: rRNA-guided normalisation

In nematodes, worm-specific argonautes (WAGOs) load sRNAs with specific properties, typically ∼22 nt in length starting with a 5’ triphosphorylated Guanosine (22G sRNAs). Immunoprecipitation and sequencing of bound sRNAs for AGOs that associate sRNAs from a broad set of loci, such as CSR-1, WAGO-4 in *C. elegans* or exWAGO in Strongylida parasites presents certain challenges. Standard normalisation methods such as TMM rely on the assumption that most loci are not differentially expressed across samples (Robinson & Oshlack, 2010). This assumption holds in experimental designs where only a subset of genes is expected to change. However, it has been previously shown that when most of the genes in a sample are differentially expressed, this assumption is invalid (Bullard et al., 2010; Y. Zhou et al., 2017), for example, in RNA-seq experiments comparing tissues with clear differences in transcriptional activity.

Here, by estimating normalisation factors based on the adjusted expression of rRNA fragments, we were able to improve the detection of genomic regions enriched in exWAGO guide production. We used rRNA because, in general, we do not expect WAGOs to associate with this type of sRNA (Seroussi et al., 2023; Zagoskin et al., 2022). Although previous reports in humans and mice have shown that rRNA fragments can be enriched in AGOs (Guan & Grigoriev, 2021), neither of our analyses in *C. elegans* WAGOs nor exWAGO show general patterns of rRNA enrichment in IPs. Other elements could be used for normalisation, for example, miRNAs could be employed for WAGOs that do not bind them. However, using rRNA allows the same approach to be used across all AGOs. This supports the use of rRNA-derived sRNAs as a stable internal reference for normalisation in this context.

Although this approach reduces the false negative rate introduced by standard TMM, we caution that this strategy is not perfect, since different amounts of rRNA fragments could originate in samples with different levels of degradation or with higher background levels. Careful inspection of the resulting MA plot is still recommended to verify that the effect of normalisation agrees with biological assumptions. In most of our samples, this normalisation worked well, but our analysis in *T. circumcincta* indicates higher levels of rRNA fragments in the IP, potentially causing an underestimation of enriched regions, leading to more conservative results than for other species. In future work, we could consider using controlled spike-ins, in both IP and input samples to evaluate alternative normalisation strategies. Nevertheless, these problems do not undermine the broader applicability of our normalisation strategy, nor the robustness of our biological conclusions, which are supported by consistent patterns of TE enrichment across Strongylida parasites.

### The dual role of exWAGO and soloLTR-derived sRNAs in Strongylida parasites

Despite their relatively low genomic span in clade V nematodes, LTR retrotransposons have been linked to key processes such as genome expansion, innate immune responses, and adaptation to environmental stress (Ansaloni et al., 2019; Kanzaki et al., 2018; Ni et al., 2016). In *C. elegans*, the argonaute protein HRDE-1 has been implicated in LTR regulation under heat shock conditions (Ni et al., 2016), and ERGO-1 has also been shown to target LTR elements (Fischer & Ruvkun, 2020). Notably, the loss of ERGO-1 in *Caenorhabditis inopinata* correlates with a marked expansion of LTR retrotransposons in that species (Kanzaki et al., 2018). Remote homology searches across predicted proteomes of Strongylida parasites revealed that ERGO-1 is absent (Supplementary Table S9). While we cannot exclude the possibility that other argonautes in Strongylida bind LTR-derived sRNAs, our enrichment analyses demonstrate that exWAGO orthologs—from *C. elegans* PPW-1 to parasitic Strongylida species—consistently associate with LTR-derived small RNAs. This suggests that exWAGO retains an ancestral role in LTR surveillance. But why does exWAGO show this remarkable preference for LTR-derived sRNAs in Strongylida parasites? An increased activity of LTR retrotransposons can potentially lead to genomic instability. In this regard, the preference for soloLTR-derived sRNAs in adult worms may represent an advantage, because by retaining transcriptional activity, they can act as a safe source for sRNAs to potentially regulate full-length elements of the same family. Although not identical, similar mechanisms are known in *Drosophila* species where genomic loci like the flamenco cluster retain soloLTRs that act as the source of piRNAs (Van Lopik et al., 2023), which in turn can modulate active LTR retrotransposons in *trans*. These parallels raise the possibility that nematodes may have independently evolved decayed LTRs as safe templates of small RNA biogenesis for TE regulation.

In addition, the immunogenic capacity we observed for *H. bakeri* EVs and the non-vesicular exWAGO (Buck et al., 2014; Neophytou et al., 2025), hints at a potential role of LTR-derived sRNAs in modulating host immunity. In this context, the preferential export of sRNAs derived from non-autonomous LTR elements may represent a mechanism by which parasites exploit genomic fossils or degenerate elements to mediate cross-species communication. Notably, although we observed that elements of undetermined conservation are consistently preferred for secretion, one limitation of our analysis is that conservation was defined based solely on the reverse transcriptase protein. Our results are consistent in that both secreted exWAGO forms show a preference for non-autonomous LTR elements. We hypothesise that these genomic fossils provide a stable, low-risk source of guides that can be safely exported without fueling retrotransposition. However, further work to identify host targets and test functional consequences of these secretion pathways will clarify how widespread and adaptive this strategy is for nematode parasitism.

Intriguingly, beyond parasitic nematodes, fungal pathogens such as *Botrytis cinerea* also rely on LTR-derived sRNAs as a source of extracellular RNA with immunomodulatory potential (Porquier et al., 2021). While *B. cinerea* preferentially uses full-length LTRs as the source of secreted RNAs, nematodes like *H. bakeri* show a preference for non-autonomous LTRs. This contrast suggests that the evolutionary paths and molecular strategies shaping extracellular RNA repertoires may differ across kingdoms, but highlight a potential convergent source of sRNAs for extracellular RNA communication.

Together, our results support a model in which exWAGO maintains genome integrity through broad TE regulation while simultaneously co-opting fragmented or non-autonomous LTR retrotransposons as platforms for regulatory sRNA production. This dual role positions exWAGO at the interface of genome defence and parasite-host communication and reveals how LTR-derived sequences can be co-opted for extracellular signalling in parasitic nematodes.

## Methods

### Transposable element annotation

We produced *de novo* repeat annotations using earlGrey v.4.0.3 (Baril et al., 2024), with the Nematoda Dfam library (Dfam 3.7), and ten iterations of the “BLAST, Extract, Extend” process. For *H. bakeri* and *H. polygyrus*, we used our previously published curated libraries (Stevens et al., 2023), with some minor updates.

Redundancy within the predicted libraries was resolved by collapsing models with at least 80 nt length and sharing 80% of nt identity in at least 80% of the model to the centroid model (the longest one), using cd-hit-est (W. Li & Godzik, 2006). To remove non-TE-related proteins, getorf from the EMBOSS package was implemented to obtain ORFs with a minimum size of 300 aa, to subsequently annotate protein domains using PfamScan (http://ftp.ebi.ac.uk/pub/databases/Pfam/Tools/) with the Pfam database (RELEASE 35). Models with hits to non-TE-related proteins were filtered out. Furthermore, TEsorter 1.4.6 was used to identify TE-related protein domains (Zhang et al., 2022). Unknown models, with hits for RT, ENDO, and INT, were further classified as LINE or LTR based on TEsorter results. Moreover, TE-Aid (available at https://github.com/clemgoub/TE-Aid) was used to identify the structural domains of predicted models. We classified them as DNA/MITE models when showing only TIR structures in self-dot plots and less than 1 kb (Goubert et al., 2022).

To have insights into the evolution of transposable elements in nematodes where exWAGO can be secreted, we manually curated the models of *N. brasiliensis* and *T. circumcincta*. For *A. ceylanicum*, we only manually curated LTR retrotransposons, and the models required to annotate 80% of the total repeat content. For this, we retrieved up to 50 instances of each element, extending 1000 nt of each edge, using the make_fasta_from_blast.sh (with the parameters 0 and 1000) and ready_for_MSA.sh (with the parameters 50 and 40) scripts. (https://github.com/annaprotasio/TE_ManAnnot). The resulting sequences were aligned using mafft v7.520, and the alignments were inspected using AliView v1.28. We identified the canonical TGT and CAC motifs at the beginning of LTRs and TAT for DNA transposons to trim divergent or poorly aligned columns (Goubert et al., 2022). For models where the edges were highly degenerate, we used CIAlign (Tumescheit et al., 2022) with the parameters:--insertion_max_size 50 --crop_ends --crop_divergent --crop_divergent_min_prop_ident.5 -- crop_divergent_min_prop_nongap.6 --crop_divergent_buffer_size 15 --make_consensus. Consensus models from the resulting alignments were built using CIAlign with the make_consensus function.

Resulting libraries were then used to produce annotations using RepeatMasker with the parameters (-s-cutoff 400-a). Since RepeatMasker produces fragmented annotations, the RepeatCraft pipeline with the Loose mode (Baril et al., 2024; Wong & Simakov, 2019) from earlGrey was used to collapse fragmented annotations. Overlapping bases between repeats were assigned using the 01.assign_TE_overlaps.R (available at https://github.com/imu93/ms_evo_exwago). Briefly, elements belonging to the same superfamily were collapsed into a single element using the reduce function from the GenomicRanges R package (Lawrence et al., 2013). If overlapping elements belonged to a different family, the family of the collapsed segment was assigned based on the longest element. Moreover, overlapping bases of elements of different super-families were assigned based on a hierarchical classification. These bases were assigned to the element that belongs to the super-family with more bases in the genome.

### Benchmark of curated TE libraries

TE library benchmarking was conducted using the get_family_summary script (available at: https://github.com/jmf422/TE_annotation/blob/master/get_family_summary_paper.sh). This script classifies consensus sequences of a tested library, based on their degree of sequence similarity, coverage and divergence, relative to a curated library. The classification consists of four categories, perfect (>95% of sequence similarity, >95% of coverage and <5% of divergence), good (same as perfect but up to <10% of divergence), present (>80% of coverage), and missed (no hits or <80% of coverage).

### ncRNA gene annotations

ncRNA annotation was carried out using several homology-based tools. We used infernal v1.1.4 and Rfam v.14.10 covariance models to annotate known ncRNA families (--rfam --cut_ga -- nohmmonly). In addition, tRNAscan-SE (-Q-E --score 40) (v1.4) and RNAmmer (-S euk-m tsu,lsu,ssu) (v1.2) were used to annotate tRNAs and rRNAs, respectively. miRNA annotation was conducted using the precursors of miRNAs for nematodes in miRBase (v.22). After obtaining precursors, blastn was used (-evalue 1e-5-perc_identity 90-qcov_hsp_perc 90). In addition, miRNA annotation of *N. brasiliensis*, *T. circumcinta* and *A. ceylanicum* was enhanced by using data from regular sRNA-seq libraries (NCBI Bioproject PRJNA1200757) using MirDeep2 (Friedländer et al., 2012).

Meanwhile, piRNAs were annotated only in a subset of species. The annotation of this ncRNA family was conducted using bowtie1 (-f-a --best --strata-v 0-m 3-S) and previously identified type 1 piRNAs in *H. bakeri*, *N. brasiliensis* (Beltran et al., 2019). Given the close phylogenetic relationship between *H. bakeri* and *H. polygyrus*, we used *H. bakeri* as a reference to annotate putative conserved piRNAs in *H. polygyrus*. Using a similar approach, we annotated piRNAs in a chromosome-level assembly of *N. brasiliensis*.

LncRNAs and lincRNAs were annotated using publicly available RNA-seq data in *H. bakeri*, *N. brasiliensis*, *T. circumcincta*, and *A. ceylanicum* (*H. bakeri*: PRJNA750155; *N. brasiliensis*: PRJEB16076, PRJEB20824, SRR25993873; *T. circumcincta*: PRJEB7677; *A. ceylanicum*: PRJNA231490). Briefly, we aligned RNA-seq reads to their respective genome using STAR v.2.7.10b using the gtf gene annotation file to define junctions. The resulting bam files were sorted using samtools sort (Li et al., 2009) to build transcriptomes with StringTie v.2.2.1 (Pertea et al., 2015). We evaluated the transcript coding potential using FEELnc_filter, cpc2, lgc, and PfamScan (Finn et al., 2014; Kang et al., 2017; G. Wang et al., 2019; Wucher et al., 2017), to filter out transcripts produced from protein-coding genes. To classify lncRNAs and lincRNAs, FEELnc_classifier was used (Wucher et al., 2017). In addition, lncRNAs and lincRNAs were annotated in *H. polygyrus* using blastn with the predicted lncRNA and lincRNAs in *H. bakeri* as reference (-evalue 1e-10-perc_identity 90 - qcov_hsp_perc 90).

### Segmented annotations

Aiming to understand the contribution of different genomic categories to sRNA production, we built non-overlapping annotations in which each base in the genome has a single annotation. We assigned overlapping regions based on a hierarchical classification using the setdiff function from the GenomicRanges R package (Lawrence et al., 2013). This annotation includes both sense and antisense annotations for each genomic region. Our hierarchy prioritises known sRNA families (in the following order: miRNA, piRNA, yRNA, rRNA, tRNA, snRNA, snoRNA, lincRNA, lncRNA), then TEs (TE families were ranked based on their genomic span), exons, introns, and finally, in the last part of the hierarchy we included unstranded repeats such as MITEs, SINEs, Low complexity repeats, Simple repeats, and Unknown repeats, in that order.

### exWAGO immunoprecipitations and RNA isolation from worms

To interrogate the sRNAs associated with exWAGO in different nematode species, including *H. bakeri*, *T. circumcincta*, *N. brasiliensis* and *A. ceylanicum* we used previously published sRNA library datasets (NCBI Bioproject PRJNA1200757) that were generated by using the eluate and unbound fraction of exWAGO immunoprecipitation from adult worms. Here we generated sRNA libraries from wild caught, naturally *H. polygyrus*-infected wood mice (*Apodemus sylvaticus*) from woodlands in Midlothian, Scotland (e.g. Penicuik Estate (55°49’N 3°15’W) and Hewan Woods, (55°52’N 3°08’W). To immunoprecipitate exWAGO from these samples, ∼1.5 cm of the duodenum was harvested and flash frozen immediately in liquid nitrogen to minimize RNA degradation, and stored in-70°C until required. Protein G beads (ThermoFisher, 10003D) were washed five times using cold Binding Wash Buffer (PBS, 0.02% Tween-20) and polyclonal rat anti-exWAGO serum antibodies (Neophytou et al., 2025) were conjugated on the prepared beads by incubation (2h, rotating wheel, 4°C). Unconjugated antibody was then removed, and the beads were equilibrated using cold lysis buffer (150 mM NaCl, 10 mM Tris.HCl, 0.5 mM EDTA, 0.5% NP40) containing protease inhibitors (1 tablet per 5 ml) (Roche, 11873580001) three times. Then, the frozen tissue was ground into a fine powder using a pestle and mortar under liquid nitrogen and then lysed in 1 ml of cold lysis buffer as above supplemented with 200 U/ml RNase inhibitors (Promega, N2515). Unlysed material was removed by centrifugation (16,100 rcf, 10 min, 4°C), and the supernatant lysate was incubated with the antibody-conjugated beads (45 min, rotating wheel, 4°C). The beads were washed with cold Low Salt buffer (50 mM Tris.HCl pH 7.5, 300 mM NaCl, 5 mM MgCl2, 0.5% NP40 and 2.5% glycerol), followed by two washes with cold High Salt buffer (50 mM Tris.HCl pH 7.5, 800 mM NaCl, 10 mM MgCl2, 0.5% NP40 and 2.5% glycerol) (5 min, rotating wheel, 4°C). The beads were then washed once with cold Low Salt buffer, followed by a cold PNK buffer wash (50 mM Tris.HCl pH 7.5, 50 mM NaCl, 10 mM MgCl2, 0.5% NP40). For small RNA sequencing analysis, the RNA was eluted directly in Qiazol (700 μl, 5 min, room temperature) (Qiagen) and stored-70°C until required.

For the adult worm *H. polygyrus* sRNA libraries, RNA was isolated from wild-derived *H. polygyrus* harvested from lab-reared wood mice at day 14 post-challenge. This outbred colony of wood mice was originally wild-caught, but is now maintained in standard laboratory conditions at the University of Edinburgh (see Sweeny et al., 2021 for more details). Worms were washed in PBS and then lysed in 700 μl of Qiazol by bead beating using 5 mm steel beads (Qiagen, 69989) using the Tissue Lyser II (Qiagen) in pre-cooled cartridges for 2 minutes at 30 Hz twice. The RNA was then processed for sequencing as described below.

### Small RNA libraries

sRNA libraries were generated as described in Neophytou et al. (2025). Briefly, RNA in Qiazol was spiked with 7 μl of 10 pM RT4 synthetic spike (CTTGCGCAGATAGTCGACACGA). The RNA was then extracted using the miRNA Serum/Plasma kit (Qiagen, 217184) and treated with RNA 5’ Polyphosphatase (Lucigen, RP8092H) according to the manufacturer’s instructions. The reaction was terminated using ethanol precipitation (-70°C, overnight), and the precipitated RNA was resuspended in 2.5 μl of nuclease-free water and 2.5 μl of Buffer 1 (TriLink, L-3206). The sRNA libraries were prepared using the CleanTag Small RNA Library Preparation Kit (TriLink, L- 3206) using half reaction volumes. The *H. polygyrus* immunoprecipitation or total adult worm samples were generated using 1:10 or 1:12 dilution of 5’ and 3’ adapters, respectively. All libraries were generated using 20 amplification cycles, and the profile of the libraries was assessed using the High Sensitivity DNA Bioanalyser chip (Agilent, 5067-4626). Pooled libraries were size-selected (140-180 bp for immunoprecipitation; 140-220 bp for total adult worms) using gel purification to remove adapter dimers. The libraries were sequenced on the Illumina NextSeq 2000 platform by the Edinburgh Clinical Research Facility using single-end 100 base pair reads.

### Small RNA: processing, alignment and quantification

In addition to the small RNA-seq libraries we produced for *H. polygyrus*, we obtained small RNA-seq libraries from NCBI SRA for *H. bakeri* (NCBI bioprojects: PRJNA481340 and PRJNA1200757)*, N. brasiliensis* (PRJNA1200757), *T. circumcincta* (PRJNA1200757), and *A. ceylanicum* (PRJNA1200757), and *C. elegans* (PRJNA860633). Illumina small RNA adapters were removed using reaper (Davis et al., 2013). FastQC and MultiQC were used to evaluate read quality (Ewels et al., 2016; Andrews & Felix, 2012). For *T. circumcincta,* we collapsed technical replicates for exWAGO IP libraries, however, due to the lack of biological replicates, technical replicates for adult total libraries were treated as independent samples. A similar approach was used for *A. ceylanicum*. Using Pullseq v.1.0.2 (available at https://github.com/bcthomas/pullseq), we then retrieved reads between 18 to 27 nucleotides for downstream analyses. ShortStack v.3.8.5 (Johnson et al., 2016) was used for genome indexing and alignment with the following parameters: bowtie-buld --offrate 2; --nohp --mincov 5 --pad 1 --mismatches 1 --mmap u -- bowtie_m all --ranmax’none’ --bowtie_cores 32 --sort_mem 248G. After using samtools split (Li et al., 2009) to obtain individual bam files, the featureCounts function from Rsubread (Liao et al., 2019) was used to quantify aligned sRNAs to segmented annotations (see Segmented annotation above) (parameters: allowMultiOverlap= TRUE, strandSpecific= 1, fracOverlap= 0.7, useMetaFeatures= TRUE, largestOverlap= TRUE).

### Differential expression and enrichment analyses of exWAGO and adult total across clade V nematodes

We performed individual differential expression analysis comparing exWAGO IP against input libraries (adult total) using edgeR (Robinson, McCarthy, and Smyth, 2010). For *C. elegans* we compared PPW-1, SAGO-1 and SAGO-2 IPs against input samples (data from Seroussi et al., 2023). We removed regions with less than five CPM in conditions with fewer replicates using the filterByExpr function from edgeR v.4.0.16 (Robinson, McCarthy, and Smyth, 2010).

We then estimated normalisation factors using the TMM normalisation method with the calcNormFactors function. After this, we fitted negative binomial generalised log-linear models using the glmFit function. After visualising diagnostic MA plots and determining that TMM assumptions are broken in these contrasts, new normalisation factors were estimated using the expression of rRNA fragments (those that mapped in sense). To account for compositional bias of rRNA in the different libraries, we used TMM on rRNA genes, reducing the effect of any extreme values for normalisation factor estimation. We then used glmFit and glmTreat to fit negative binomial generalised log-linear models and determine enriched regions. Genomic regions that were not enriched in the IPs were considered as unbound.

### Argonaute sRNA guide diversity in Strongylida and C. elegans

To compare the diversity of sRNA guides between exWAGO orthologs in Strongylida parasites and *C. elegans* argonautes, aiming to account for differences in sequencing depth between IP experiments, we normalized raw counts into counts per million (CPM). We then used the vegan R package to estimate Shannon’s and Simpson’s diversity indices (Oksanen et al., 2012). Statistical comparison was then carried out using the ggpubr R package (Kassambara, 2025).

### Structural LTR annotation, sRNA quantification and differential expression analysis

To classify the structural features of the initial genomic regions annotated as LTR retrotransposons, we first split the curated consensus sequences of LTR retrotransposons into the LTR (directed repeats) and internal region regions (containing CDS of LTR proteins), using selfblast produced with TE-Aid, and an *ad hoc* R script (available at: https://github.com/imu93/ms_evo_exwago). Using the split models, we build a new LTR library. After splitting the models, we selected only one LTR region per consensus model. In models where the nucleotide identity was lower than 80% on their directed repeats or lacked directed repeats, we maintain the model, but without structural information. Using RepeatMasker (-s-cutoff 400-a-nlow) (Smit et al., 2013), we then reannotate all the genomic segments previously identified as LTR retrotransposons. These new annotations were used to quantify the assigned sRNAs using the featureCounts function from the Rsubread package (strandSpecific= 1, fracOverlap= 1, useMetaFeatures= TRUE, largestOverlap= TRUE) (Liao et al., 2019).

We further performed differential expression analysis using edgeR v.4.0.16 (Robinson, McCarthy, and Smyth, 2010). First, we performed differential expressions analysis between IP against Unbound for both vesicular and non-vesicular exWAGO, and adult IP against adult total, using housekeeping normalisation with rRNA as a reference category. We then fit negative binomial generalised log-linear models with glmFit, and define differentially expressed regions using glmTreat (FDR < 0.05 and log_2_ FC = 1 for secreted exWAGO contrasts and FDR < 0.01 and log_2_ FC = 1 for adult IP against adult total). Finally, using the union of differentially expressed regions, we performed a secreted IP against adult IP contrast. Since this contrast does not involve the comparison of unbalanced conditions, we used TMM for normalisation factor estimation, followed by glmFit and glmTreat (FDR < 0.05 and log_2_ FC = log_2_(1.2)).

### Orthology inference of LTR retrotransposons

We performed orthology inference of LTR retrotransposon families among clade V nematodes based on protein identification and phylogenetic reconstruction using Orthofinder v.2.5.4 (Emms and Kelly, 2019). For this, we first annotated LTR proteins on genomic copies using TEsorter (Zhang et al., 2022) with Hidden Markov Models (HMMs) from the metazoan Rex database. We then used an *ad hoc* R script to select TE LTR proteins with at least 80% of model coverage and E-value < 1e-5.

The orthology of predicted protein domains was interrogated using Orthofinder (Emms and Kelly, 2019) to produce phylogenetic hierarchical orthogroups and build phylogenetic trees. For clustering, an inflation value of 6 was used. In addition, a phylogenetic tree based on 443 single-copy ortholog BUSCO genes was used to reconcile gene trees.

### Generalised linear mixed model for LTR-derived sRNA production

Using expression values, conservation, divergence and structural information from LTR retrotransposons enriched in exWAGO guide production, we fitted a generalized linear mixed model using the glmmTMB R package (McGillycuddy et al., 2025). We estimate log2 reads per kilobase per million (RPKM) values using the rpkm function from edgeR (Robinson, McCarthy, and Smyth, 2010), we then mean-centred the values by subtracting the mean log2 RPKM estimated for antisense-derived sRNAs. As fixed categorical effects, we used conservation encoded as highly conserved (HC), conserved (C), lowly conserved (LC) and undetermined. In addition, we modelled LTR type as autonomous (considering the presence of GAG, protease, endonuclease, reverse transcriptase, endonuclease, and integrase), non-autonomous (lacking at least one protein) and soloLTR (based on the LTR structure, lack of coding potential and surrounding LTR segments). We also used the RepeatMasker estimated relative divergence values per copy as a continuous fixed effect (Smit et al., 2013). To account for the variance each family has, we also modelled family as a categorical random effect. The same model was independently used for each species, and per-species model fit was assessed using diagnostic qq, residual vs fitted, residual density and fixed effect distribution plots, with no major deviations from normality (Fig. S9).

### Binomial generalised linear model for secreted exWAGO guides

To assess which factors influence exWAGO guide secretion from LTR retrotransposons, we fitted a binomial generalised linear model using the lme4 R package (Bates et al., 2015). We fitted enrichment in exWAGO EV IP and non-enrichment as a dichotomic response variable. As explanatory variables, we fitted conservation (based on previously defined phylogenetic hierarchical orthogroups HOGs), strand (sense or antisense), and structure (soloLTR, LTR region, internal-coding region) as categorical. In addition, to account for differences in length, we fitted log_10_-transformed mean-centred length as continuous. Model assumptions were inspected using the DHARMa R package (Hartig, 2024), with no deviations observed in the simulated residuals (Fig. S11).

## Acknowledgments

We acknowledge Jennifer Mcintyre, for the prepublication access to the *Teladorsagia circumcincta* genome assembly and genome annotation. We acknowledge Clément Goubert for his valuable insight into the transposon manual curation. Computation was performed on the Ashworth Compute Co-operative Cluster (AC3).

## Data Availability

New small RNA-seq libraries for exWAGO IPs and total adults, from *H. polygyrus* are available at the NCBI SRA under BioProject PRJNA1327922. Previously published sRNA-Seq data that were reanalysed in this manuscript are available under BioProjects PRJNA1200757 and PRJNA481340. Scripts related to differential expression analysis, genome annotation, and repeat libraries are available at https://github.com/imu93/ms_evo_exwago. Curated repeat libraries were also submitted to Dfam.

