## Supplementary Figures for "Comparative transcriptomics reveals an extracellular worm argonaute as an ancestral regulator of LTR retrotransposons"

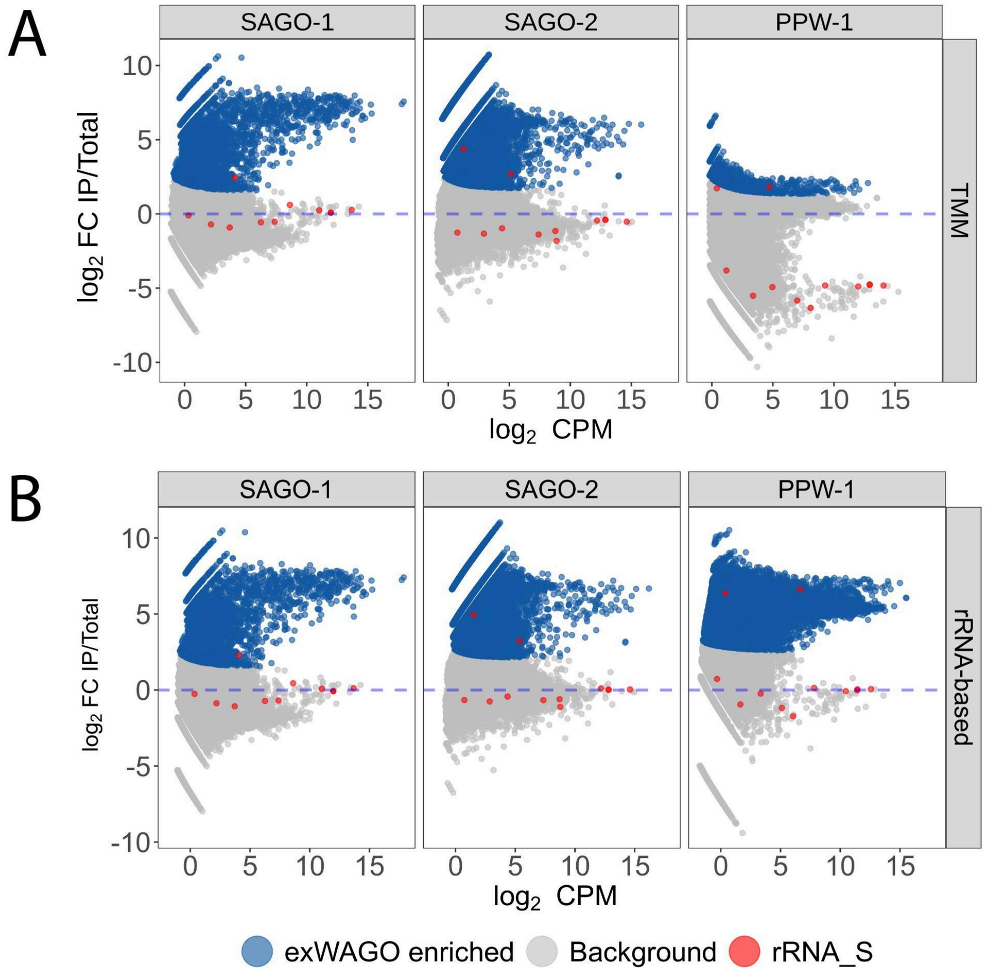


**Figure S1. Differential expression analysis for *C.elegans* SAGO-1, SAGO-2 and PPW-1 IPs, using TMM (A) or rRNA-based (B) normalisation factors.** Each dot in the MA plots represents a non-overlapping genomic region. Blue dots highlight regions significantly enriched in IP, with remaining dots in grey, suggesting regions producing mostly unbound sRNAs. Horizontal dotted lines indicate logFC = zero. Red dots indicate the expression of rRNA.

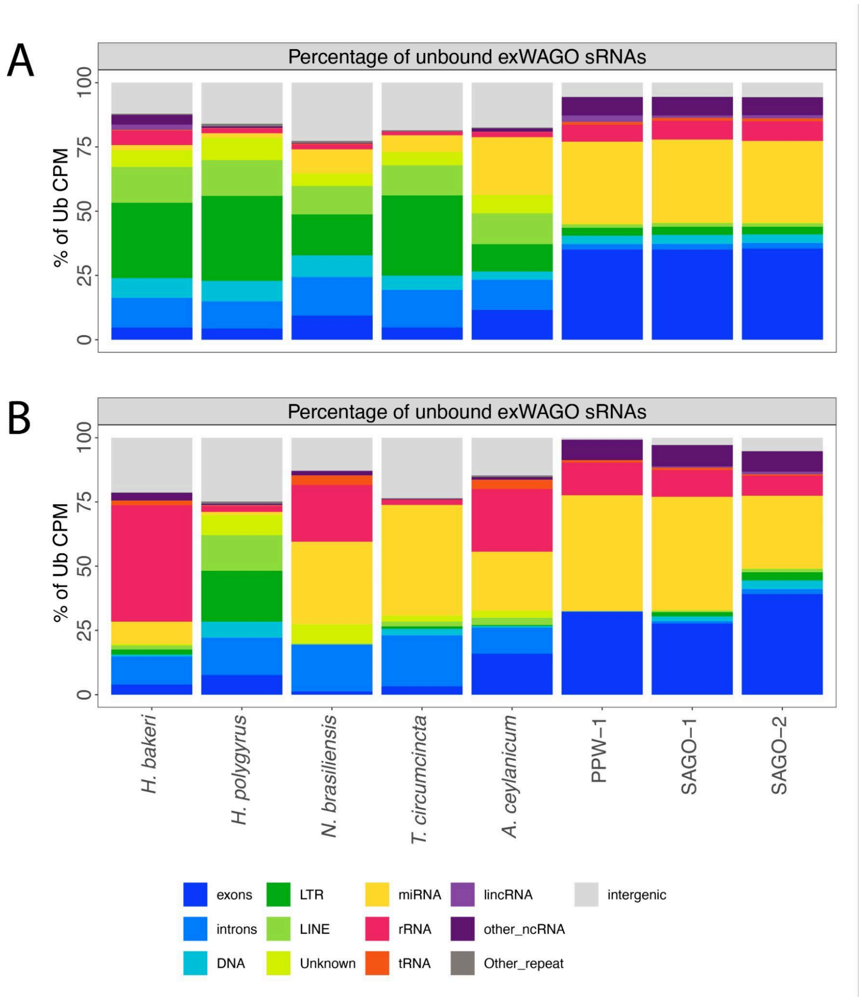


**Figure S2. Comparison of sRNAs predicted to remain unbound to exWAGO and its orthologs in Strongylida and *C. elegans*.** The distribution of the percentage of counts per million (CPM) for unbound friends using trimmed mean of M-values (TMM) is shown in (A), and the same for rRNA-based housekeeping normalisation in (B). Bars were coloured according to the annotated genomic regions.

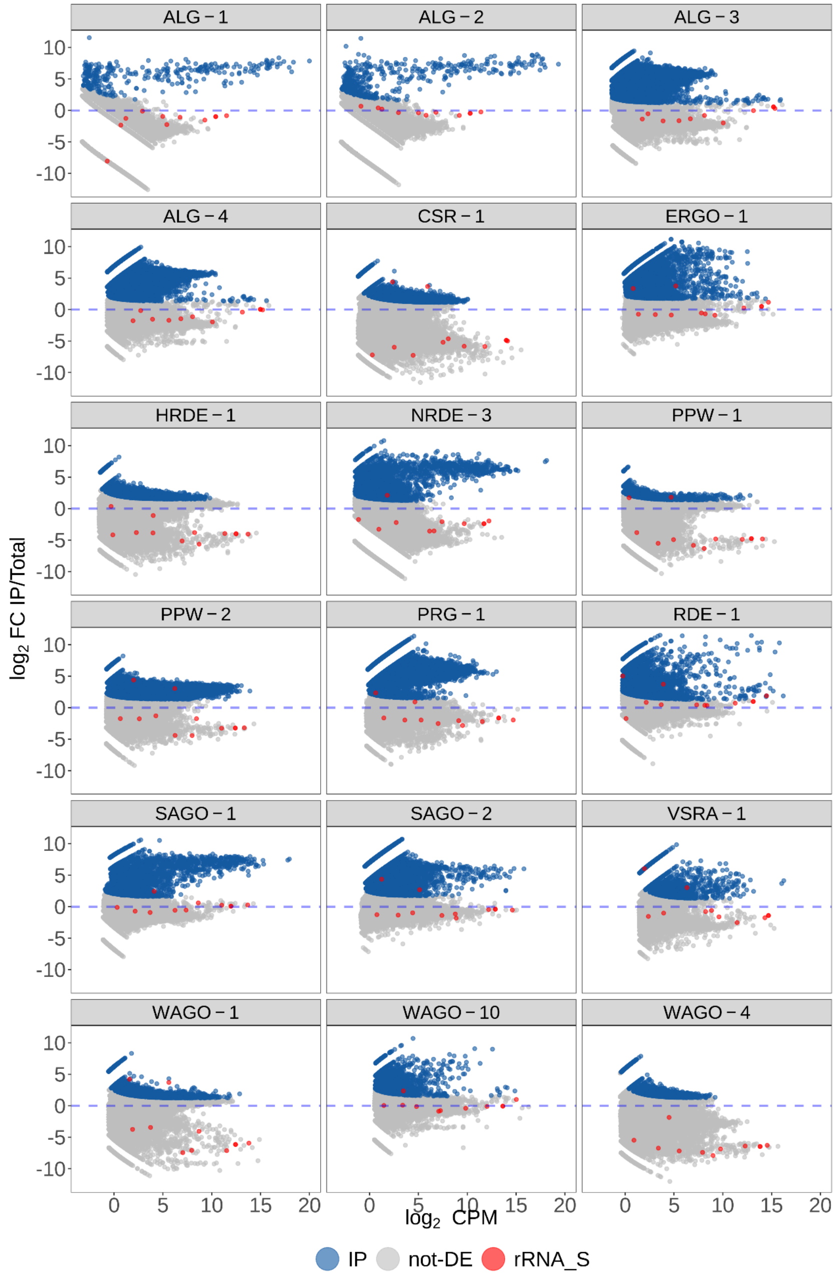


**Figure S3. Differential expression analyses using trimmed mean of M-values (TMM) of non-miRNA-related *C. elegans* argonaute IPs against input.** Blue dots on each MA plot show the genomic regions enriched in IP, grey dots represent unbound genomic regions, and red dots represent expression values for sense-stranded rRNA fragments (FDR < 0.05).

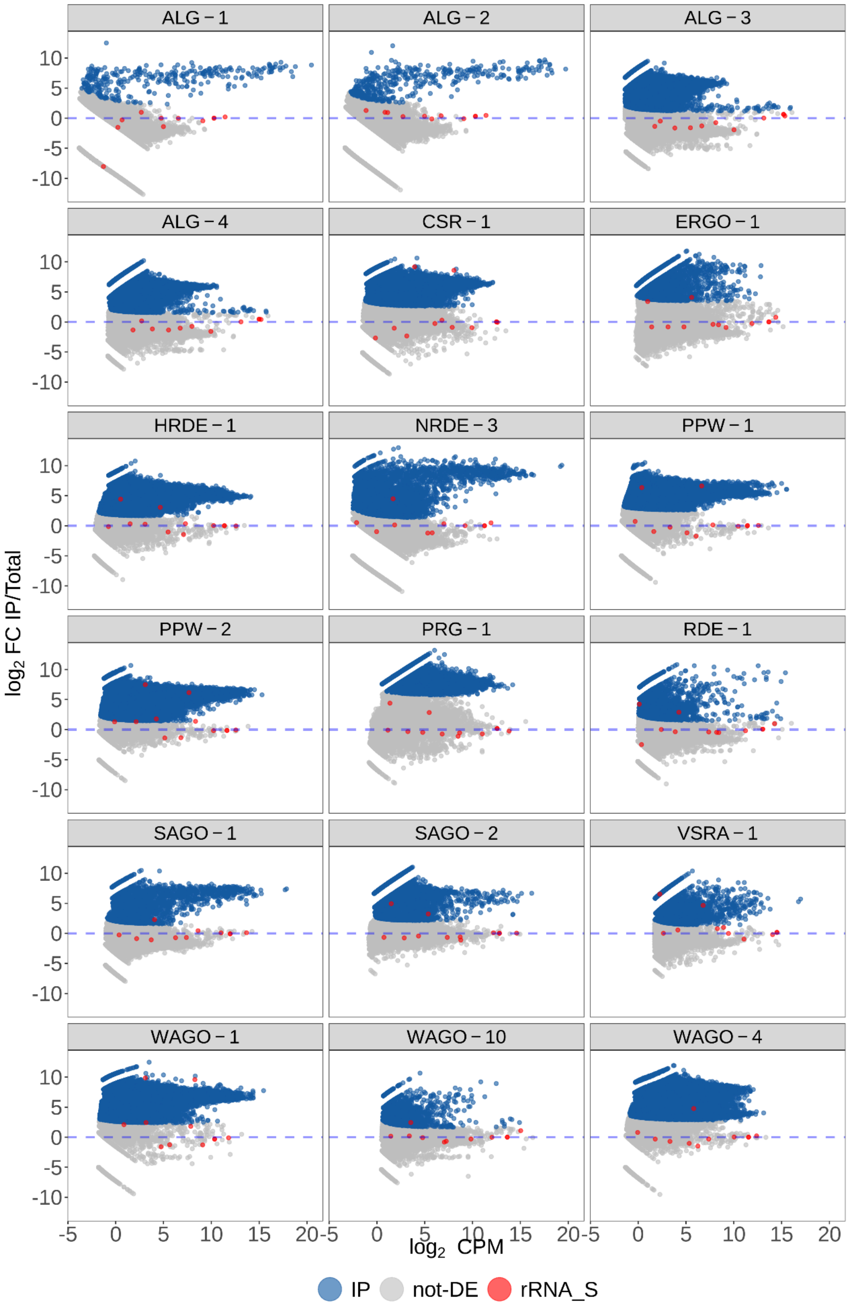


**Figure S4. Differential expression analyses using miRNA-based housekeeping normalisation of non-miRNA-related *C. elegans* argonaute IPs against input.** Blue dots on each MA plot show the genomic regions enriched in IP, grey dots represent unbound genomic regions, and red dots represent expression values for sense-stranded rRNA fragments (Contrast-specific cutoff values were used in this analysis) (see Supplementary Table).

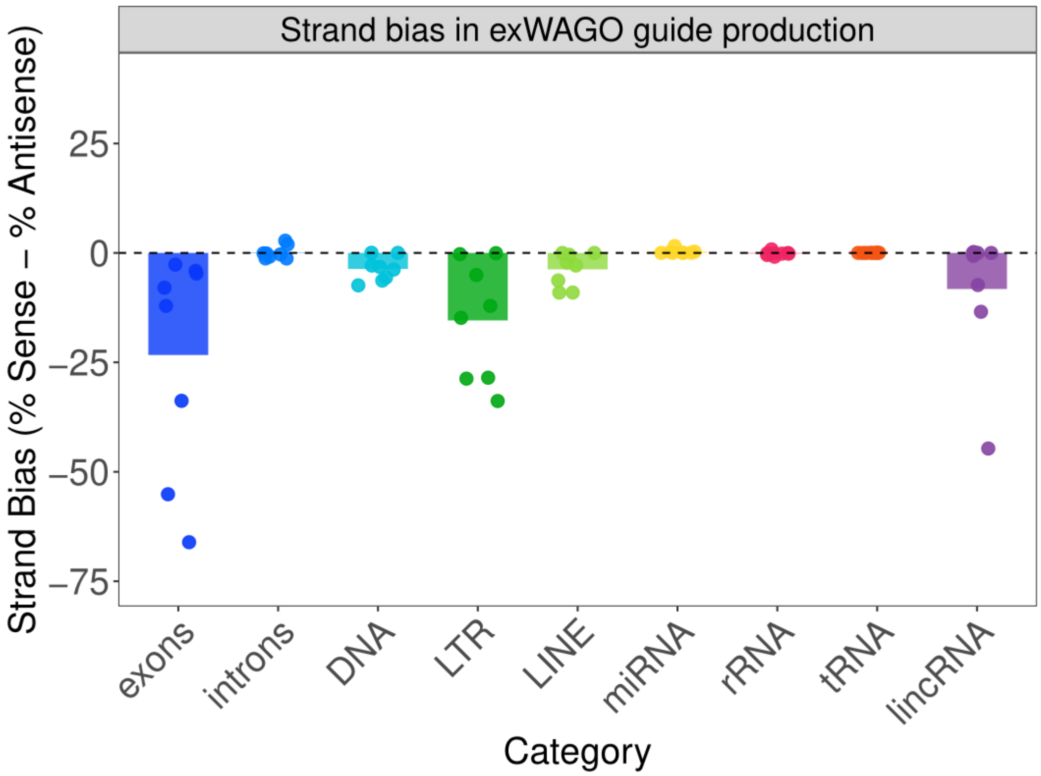


**Figure S5. Antisense strand bias in exWAGO and *C. elegans* orthologs guides.** Individual dots represent strand bias values calculated per genomic category and per species, for regions enriched in sRNA guide production. Each bar represents the mean strand bias estimated for all the species. A value of 0 (black dashed line) indicates equal sense and antisense guide production. Positive values indicate a bias toward sense-strand guides, while negative values indicate a bias toward antisense-strand guides.

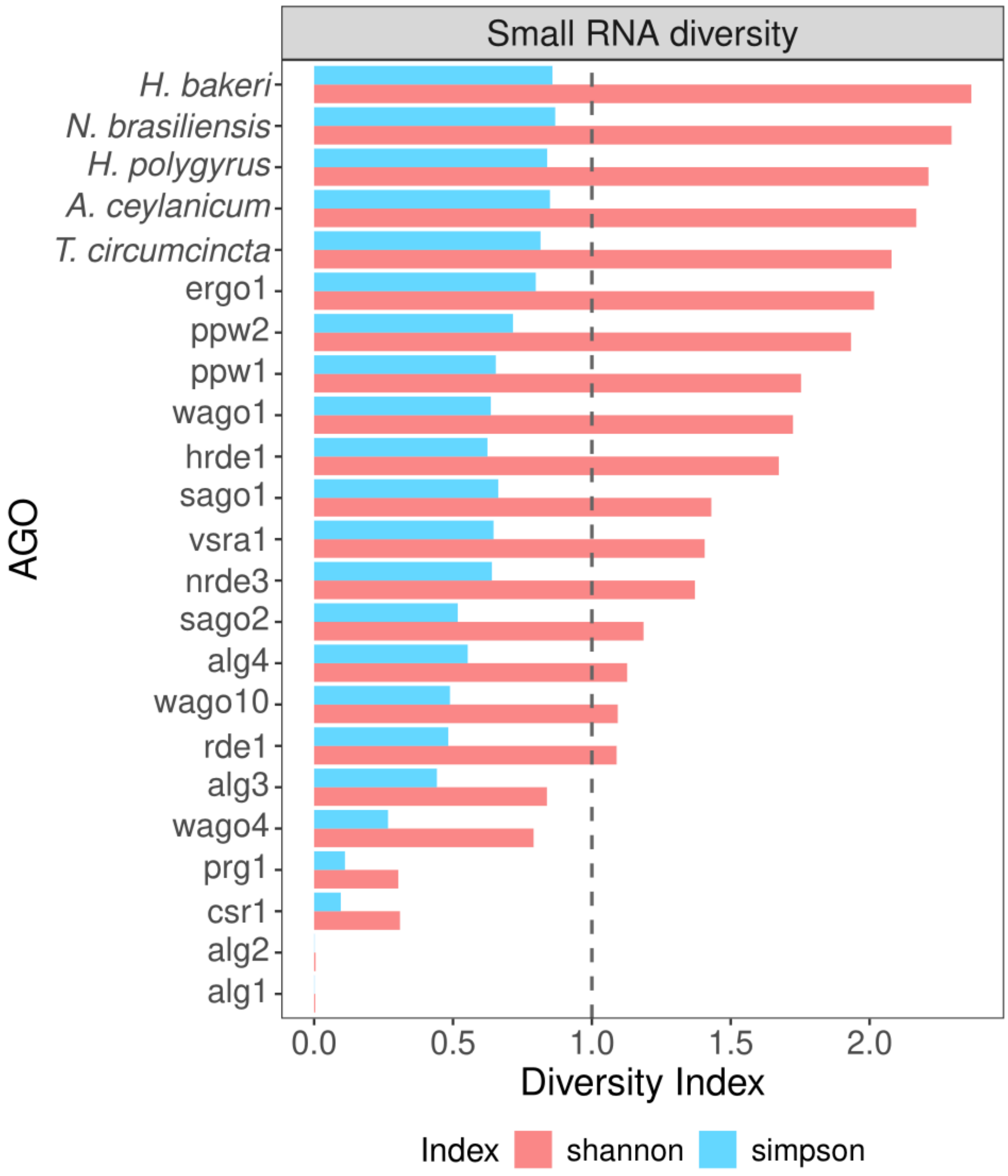


**Figure S6. Small RNA diversity in *C. elegans* AGOs and Strongylida parasites.** Small RNA diversity represented by Shanon and Simpson indices in *C. elegans* AGOs and exWAGO in Strongylida parasites. The grey dotted line indicates the maximum value (1) of the Simpson index.

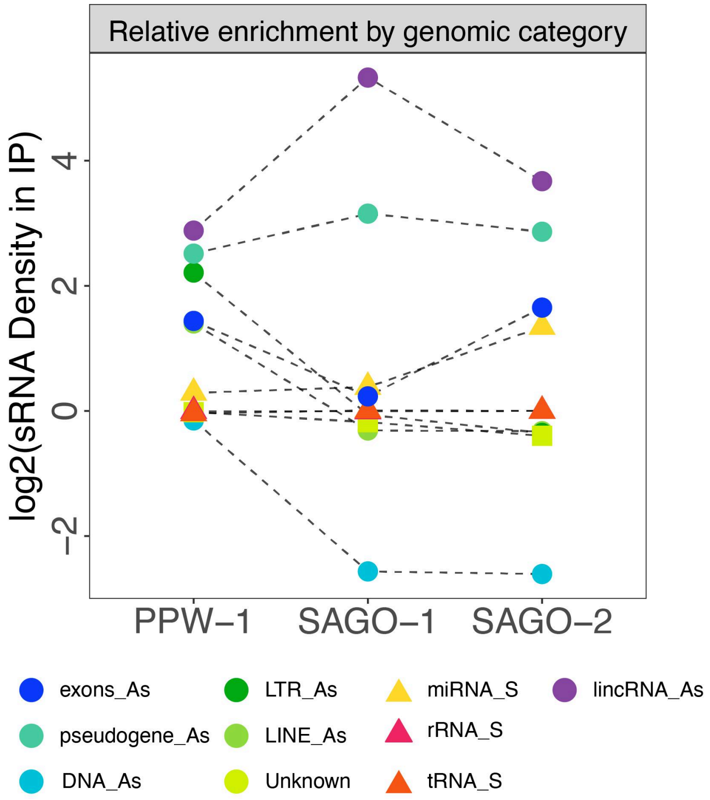


**Figure S7. Relative enrichment by genomic categories in exWAGO orthologs in *C. elegans*.** sRNA density was estimated using the percentage of Counts per Million (CPM) of enriched regions after differential expression analysis against input, divided by the genomic percentage per genomic category (log2-transformed; a pseudocount of 1 was added to both the numerator and denominator). The three exWAGO orthologs in *C. elegans* show consistent enrichment for antisense-derived lincRNAs and pseudogenes (considering only their respective exons).

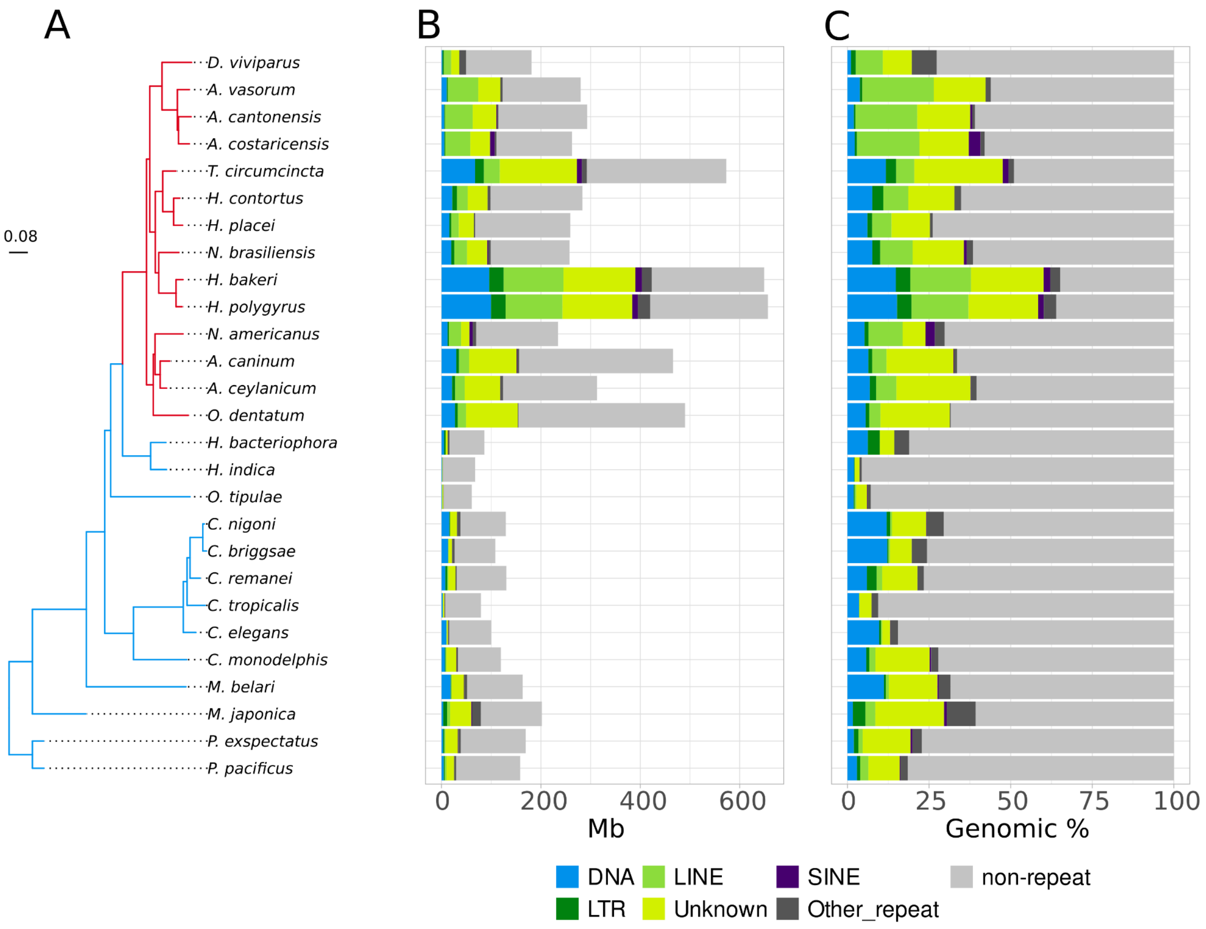


**Figure S8. Repeat landscape in clade V nematodes.** Repeat annotation after removing overlaps between elements of the same class. Maximum likelihood phylogeny inferred with a concatenated alignment of 443 complete BUSCO proteins present in the 27 nematode genomes. Red and Blue branches represent nematode species within the Strongylida and Rhabditida orders, respectively. The tree scale bar corresponds to 0.08 amino acid substitutions per site (A). Repeat span by class is shown in Mb. The category classified as non-repeat (see legend) represents all the genomic bases that were not annotated as TE or repeat. The Other repeat category represents the genomic bases annotated as Simple repeat or Low complexity repeat. RC/Helitron and DNA/MITE were collapsed within the DNA class (B). Repeat span by class as a genomic proportion (C).

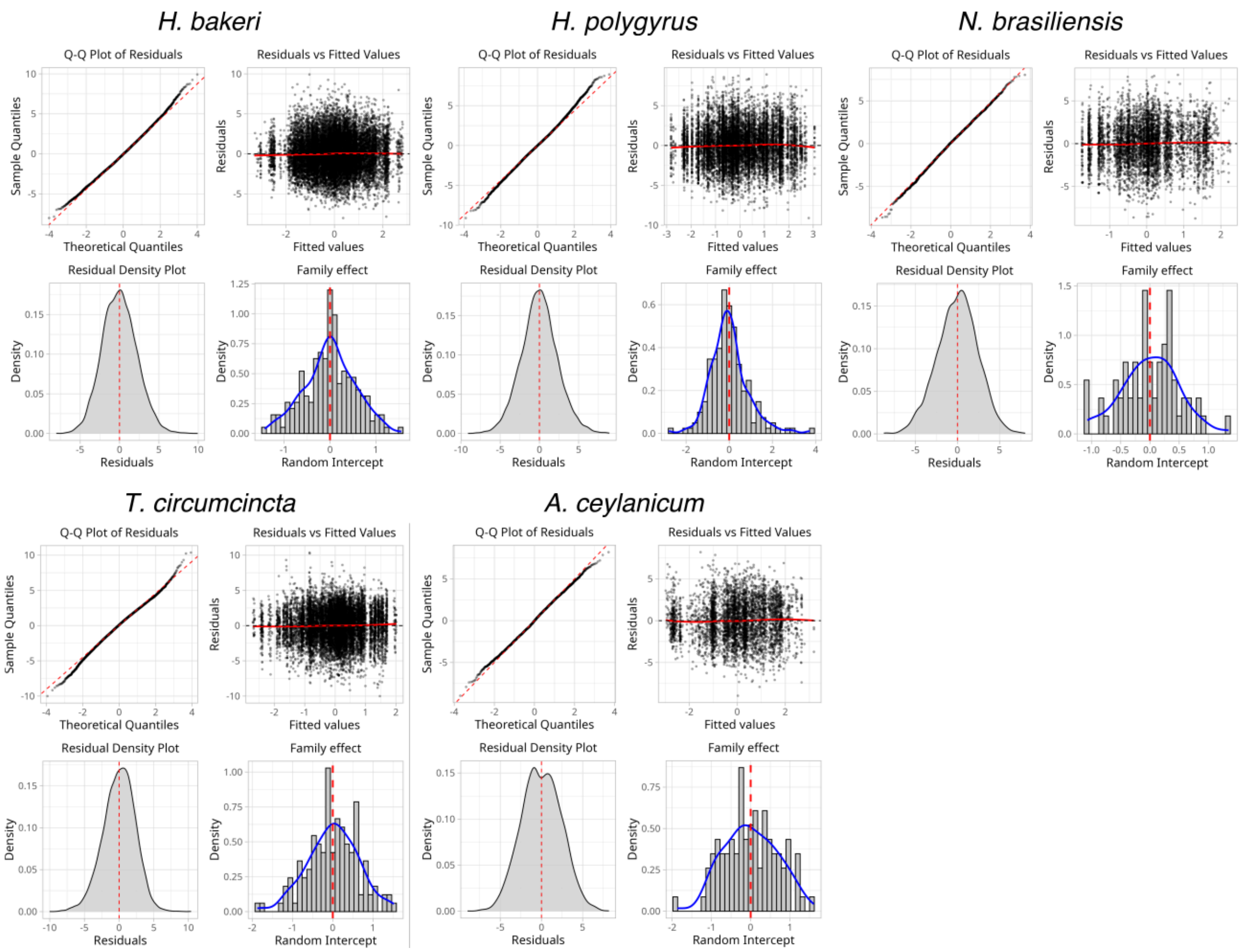


**Fig S9. Generalised linear mixed model diagnostics for LTR-derived exWAGO guide production in Strongylida.** Each panel shows diagnostic plots evaluating the fit of GLMMs, for LTR retrotransposons and modelled log2-centred expression of antisense sRNAs as the response variable. (1) Q–Q plot of residuals with theoretical normal distribution; (2) residuals vs. fitted values; (3) residual density plot; and (4) distribution of random intercepts for transposon family.

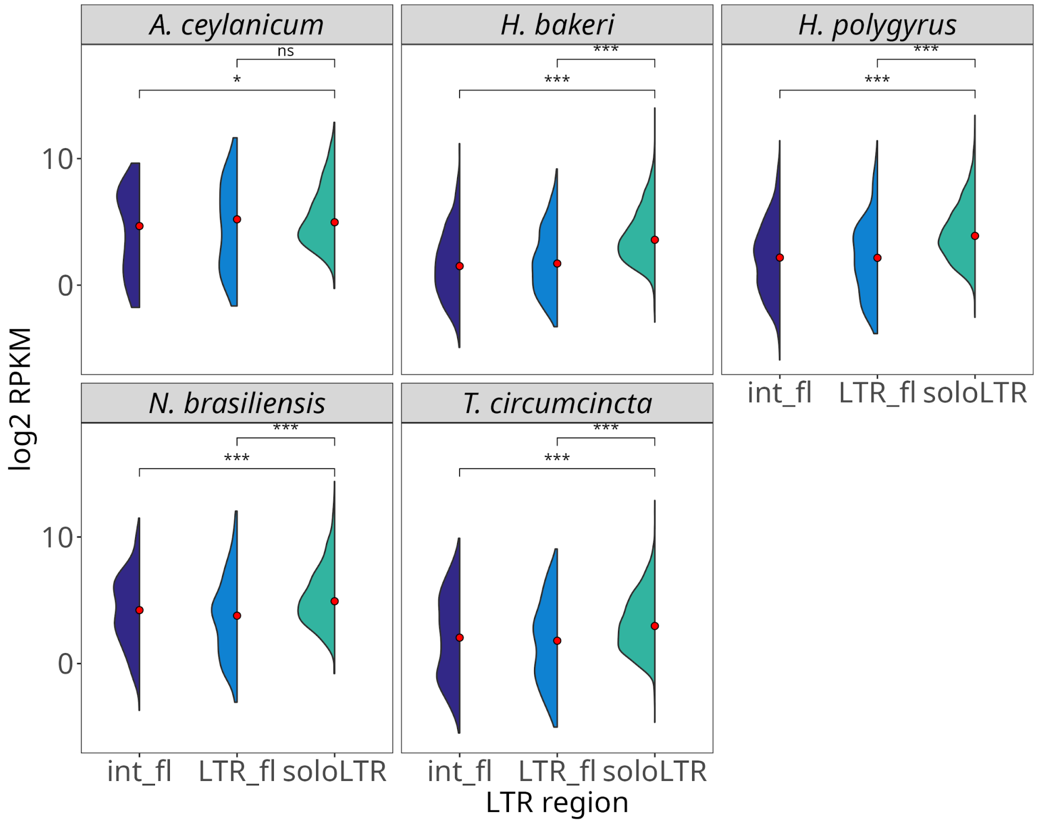


**Fig. S10. Comparison of exWAGO guide production between full-length LTRs, considering their internal (int_fl) and LTR (LTR_fl) regions against soloLTRs in Strongyloda parasites.** The x-axis shows the distribution per category, and the y-axis shows log2 reads per kilobase per million (RPKM), for LTRs enriched in exWAGO guide production. The red dots inside each distribution represent the median value per category. A one-tailed Wilcoxon Rank Sum test was used for comparisons (ns not-significant; * p-value < 0.05).

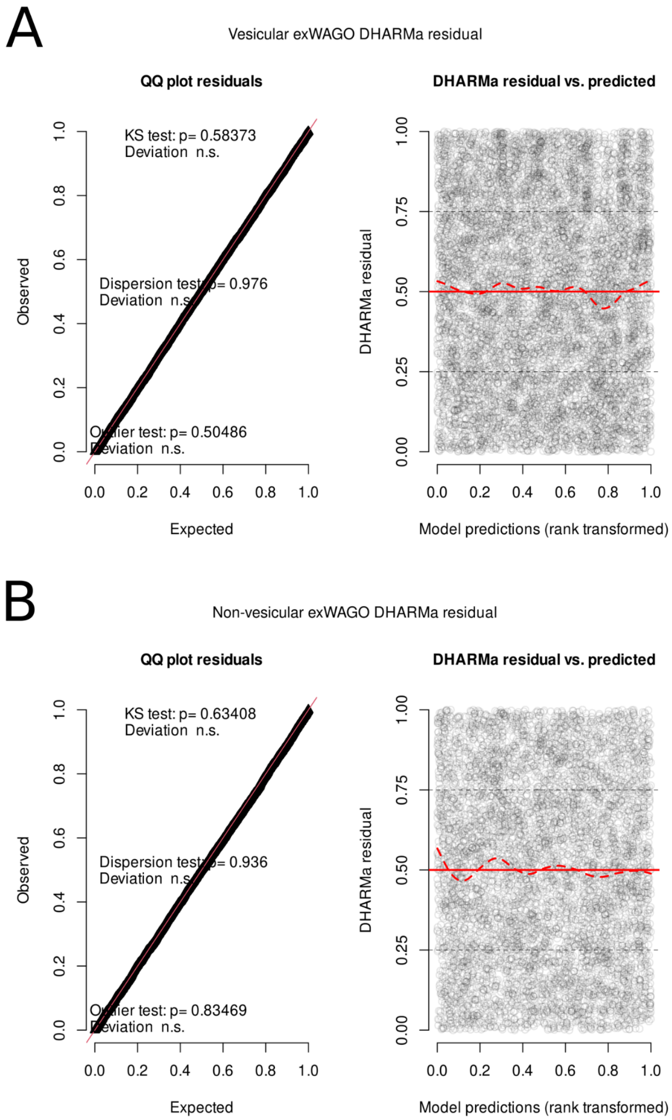


**Fig. S11. Diagnostic residual plots for the binomial generalised linear model assessing secretion of exWAGO-bound LTR-derived sRNAs.** Q–Q plots of simulated scaled residuals against the expected uniform distribution, and residual vs. predicted values are shown for vesicular exWAGO (A) and non-vesicular exWAGO (B). Kolmogorov–Smirnov goodness-of-fit test show no significant deviations from the theoretical distribution ($p > .05$ in both exWAGO forms). Dispersion and outlier tests indicate no overdispersion or outlier inflation ($p > 0.5$). The red line indicates the 1:1. Residuals vs. predicted values, showing no systematic pattern or evidence of misfit. The dashed red line represents a smoothed LOESS fit of the residuals, while the solid red line shows the expected mean residual value.
